# IL-11 disrupts alveolar epithelial progenitor function

**DOI:** 10.1101/2022.11.11.516088

**Authors:** Rosa K. Kortekaas, Kerstin E. Geillinger-Kästle, Theo Borghuis, Kaoutar Belharch, Megan Webster, Wim Timens, Janette K. Burgess, Reinoud Gosens

**Author notes:** corresponding author, Department of Molecular Pharmacology, University of Groningen, Antonius Deusinglaan 1, 9713 AV Groningen, the Netherlands. current affiliation: Newcells Biotech, Newcastle upon Tyne, United Kingdom (Megan Webster). authors contributed equally (Janette K. Burgess, Reinoud Gosens).

## Abstract

IL-11 is linked to the pathogenesis of idiopathic pulmonary fibrosis (IPF), since IL-11 induces myofibroblast differentiation and stimulates their excessive collagen deposition in the lung. The alveolar architecture is disrupted in IPF, yet the effect of IL-11 on dysregulated alveolar repair associated with IPF remains to be elucidated.

We hypothesized that epithelial-fibroblast communication associated with lung repair is disrupted by IL-11. Thus, we studied whether IL-11 affects the repair responses of alveolar lung epithelium using mouse lung organoids and precision cut lung slices (PCLS). Additionally, we assessed the anatomical distribution of IL-11 and IL-11 receptor in human control and IPF lungs using immunohistochemistry.

IL-11 protein was observed in human control lungs in airway epithelium, macrophages and in IPF lungs, in areas of AT2 cell hyperplasia. IL-11R staining was predominantly present in smooth muscle and macrophages. In mouse organoid co-cultures of epithelial cells with lung fibroblasts, IL-11 decreased organoid number and reduced the fraction of pro-SPC expressing organoids, indicating dysfunctional regeneration initiated by epithelial progenitors. In mouse PCLS alveolar marker gene expression declined, whereas airway markers were increased. The response of primary human fibroblasts to IL-11 on gene expression level was minimal, though bulk RNA-sequencing revealed IL-11 modulated a number of processes which may play a role in IPF, including unfolded protein response, glycolysis and Notch signaling.

In conclusion, IL-11 disrupts alveolar epithelial regeneration by inhibiting progenitor activation and suppressing the formation of mature alveolar epithelial cells. The contribution of dysregulated fibroblast – epithelial communication to this process appears to be limited.

## Introduction

Idiopathic pulmonary fibrosis (IPF) is a chronic progressive lung disease, which can develop as a consequence of exposure to environmental risk factors in combination with genetic predisposition. It is believed to be induced by repetitive micro-injuries to the alveolar epithelium leading to defective epithelial-fibroblast communication and chronic activation of extracellular matrix (ECM)-producing myofibroblasts resulting in excessive deposition of ECM and destruction of the physiological alveolar structure. This causes disrupted gas exchange and abnormal compliance and lung stiffness, and ultimately, respiratory failure. The median survival time after diagnosis of IPF is only 2-4 years [1]. While the recently approved drugs nintedanib and pirfenidone both reduce lung function decline and likely prolong survival, they are not able to prevent progression of established fibrosis [2–4]. Therefore, it remains essential to study new therapeutic strategies for IPF.

IL-11 is a member of the IL-6 family, which can affect various cellular processes including hematopoiesis, pro-inflammatory cytokine release by macrophages, T helper (Th) cell polarization, intestinal epithelial cell proliferation and hepatocyte apoptosis [5–9]. Interestingly, IL-11 protein levels are commonly not or hardly detectable in healthy individuals [10–12], and IL-11Rα deficient individuals are reported to be almost completely healthy, though they may suffer from the skull disorder craniosynostosis [13]. This suggests that IL-11 is not essential for normal homeostasis, whereas it is associated with various pathological conditions, including cancer and fibrosis in numerous organs [14,15]. Although fibroblasts and epithelial cells have been proposed as predominant sources of IL-11 in the lung [12,16], it is not yet established which cell types primarily produce IL-11 and show IL-11 receptor (IL-11R) expression in the human lung. While IL-11 was found to inhibit fibrotic scarring in zebrafish [17], recent mouse and human studies indicated that IL-11 may play a fundamental role in IPF pathogenesis. IL-11 gene expression is upregulated in IPF lungs, and fibroblasts isolated from IPF tissue secrete higher amounts of IL-11 protein than controls, both at baseline and in response to transforming growth factor β (TGFβ). IL-11 can induce fibroblast to myofibroblast transition, stimulate collagen secretion by myofibroblasts, and the fibrotic response in a bleomycin mouse model was diminished in fibroblast-specific IL-11Rα knock-out mice and by the administration of an IL-11 antibody in wild type mice [12].

Although the alveolar architecture is also severely affected in IPF, the effect of IL-11 on the alveolar epithelium has only been studied to a limited extent [16,18]. Yet, dysfunctional alveolar type 2 cells (AT2 cells) are believed to play a fundamental role in the pathogenesis of IPF, contributing to the initiation of fibrosis [19,20]. Furthermore, AT2 cells function as progenitor cells through their ability to trans-differentiate into AT1 cells [21]. In IPF, the regenerative capacity of AT2 cells is impaired [20], while loss of both AT1 and AT2 cells have been observed in IPF tissue. Additionally, AT2 cells in IPF were reported to show abnormalities such as endoplasmic reticulum (ER) stress, mitochondrial dysfunction, senescence and increased release of pro-fibrotic factors [1,22].

Since IL-11 was found to influence IPF pathogenesis by acting on fibroblasts [12], we hypothesized it also negatively impacts epithelial-fibroblast communication associated with normal lung repair. Thus, we studied whether IL-11 affects the repair responses of the alveolar epithelium. We first explored the protein expression pattern of IL-11 and its receptor in human lung tissue to gain insight in the predominant producer and responder cells of IL-11, and assessed differences in expression between histologically normal control lung tissue and IPF tissue. We then studied the influence of IL-11 on epithelial progenitor cell function using a mouse organoid model. Finally, we examined whether IL-11 induced dysfunctional fibroblast-epithelial communication, thereby affecting progenitor cell behavior.

## Methods

### Animal handling

Pathogen-free wild type C57BL/6J mice (>8 weeks of age, both male and female) were used in this study. Animals were housed under a 12 hour light/dark cycle with controlled humidity at room temperature (24 ± 1 °C). Water and food were provided ad libitum. All experiments were performed according to the national guidelines and upon approval of the experimental procedures by the local Animal Care and Use committee of the University of Groningen.

### Immunohistochemistry

The immunohistochemical stainings performed in this manuscript were conducted as part of the HOLLAND (HistopathOLogy of Lung Aging aNd COPD) project executed at the department of Pathology and Medical Biology of the University Medical Center Groningen, based on established protocols [23]. Lung tissue was derived from left-over lung tissue derived from lung resection and lung transplant surgeries. The study protocol was consistent with the Research Code of the University Medical Center Groningen (https://umcgresearch.org/nl/w/research-code-umcg), as well as Dutch national guidelines on ethics and professionalism (https://www.coreon.org/). Lung tissue from tumor resection surgery was taken as far away as possible from the tumor site and all tissues were checked by an experienced pathologist to ensure no abnormalities were present. For this manuscript histologically normal control lung tissue (hereafter termed ‘control’) and tissue from individuals with IPF was included. All donors were non-smokers, ex-smokers or current smokers, with an age range between 37 and 67 years.

Paraffin-embedded human lung tissue of control and IPF donors was cut with a Microm HM355S microtome (Thermo Fisher Scientific, Waltham, MA, USA) into serial sections of 3 μm. Clinical characteristics for these donors are described in table 1; we included the best FEV1/FVC value that was available from either the pre or post BD measurements. Immunohistochemistry for the detection of either IL-11 or IL-11R was performed in one batch for all sections. Tissue sections were deparaffinised in xylene and rehydrated through a serial dilution of ethanol. Heat-induced epitope retrieval was performed in a Pascall S2800 DakoCytomation pressure cooker using 0.1M Tris/HCl buffer pH=9.0. Sections were washed in PBS and incubated overnight at 4°C with primary antibodies rabbit anti-IL-11 (LSBio, Seattle, WA, USA, #LS-B15705) or rabbit anti-IL-11R (Abcam, Cambridge, UK, #ab125015, RRID: AB_109750018), diluted 1:100 (final concentration 17.1 μg/ml) or 1:400, respectively, in 1% (w/v) BSA 2% donkey serum (Jackson Immunoresearch, West Grove, PA, USA, #017-000-121, RRID: AB_2337258) in PBS. Sections were then washed with PBS and incubated with 0.3% H2O2 in PBS for 30 min. Subsequently, sections were incubated with the secondary antibody peroxidase-conjugated donkey anti-rabbit (Jackson Immunoresearch, #711-035-152, RRID: AB_10015282) diluted 1:500 in 1% BSA 2% donkey serum in PBS for 30 min at room temperature. Finally, the immunohistology reaction product was developed with NovaRED (Vector Laboratories, Burlingame, CA, USA #SK-4800, RRID: AB_2336845) for 10 min, followed by hematoxylin counterstain. They were then dehydrated through a series of ethanol-xylene and covered with xylene and Tissue-Tek^®^ coverslipping film (Sakura Finetek, Tokyo, Japan, #4770). Negative control slides were included which were handled equally, except no primary antibody was added. Slides were scanned with a Hamamatsu NanoZoomer 2.0HT digital slide scanner (Hamamatsu Photonic K.K., Hamamatsu City, Japan) at magnification 40x. Aperio ImageScope V.12.4.3 (Leica Biosystems, Wetzlar, Germany) was used to view digital sections and to obtain descriptive images. For quantitative analysis, artefacts such as tar, air bubbles and fibers were removed from images, which were otherwise unaltered. Per immunohistochemical staining, representative images of weak to strong staining were used to set a color deconvolution vector which adequately separated the blue (hematoxylin) and red (NovaRED) channel, and to set a threshold for both the blue (hematoxylin) and red (NovaRED) channels separately. Fiji/ImageJ software (ImageJ 2.1.0) [24] was used to quantify the mean intensity and percentage of positive stained area for each protein. Data analyses were performed using R software V.4.0.0 (Boston, Massachusetts, USA) using the following formulas:

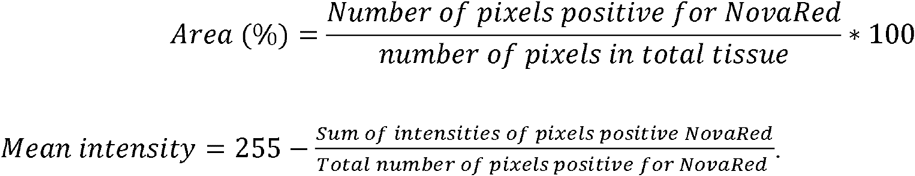

**Table 1:** patient characteristics of human lung tissue samples stained for IL-11 and IL-11R

| Patient group | Sex<br>M/F | Age<br>Median | Smoking status<br>Never/ex/current | Pack years<br>Median (range) | FEV/FVC (best)<br>Median (range) | FEV1 (% of predicted pre BD) |
| --- | --- | --- | --- | --- | --- | --- |

|  |  | (range) |  |  |  | Median (range) |
| --- | --- | --- | --- | --- | --- | --- |
| Control | 3/6 | 57 (43-62) | 0/9/0 | Ex: 15 (1.5-25)<br>3 unknown | 76 (69-86.6) | 103.6 (85-127)<br>1 unknown |
| IPF | 9/3 | 57 (37-67) | 5/6/1 | Ex: 26 (6-40)<br>1 unknown<br>Current:<br>unknown | 87.9 (67.5-100) | 49.4 (18-74.5)<br>4 unknown |

### Fibroblast isolation and cell culture

Primary human lung fibroblasts of control and IPF donors were isolated from resected tissue and from lungs that were excised during organ transplantation respectively. Lung tissue was cut into blocks of 1 mm^3^ and 2 blocks were transferred into 1 well of a 12-well plate, allowed to attach and cultured with 1 ml Ham’s F12 with 10% (v/v) fetal bovine serum (FBS, PAA Laboratories, Pasching, Austria), 100 U/ml penicillin/streptomycin (Gibco under Thermo Fisher Scientific, Waltham, MA, USA, #15070-063) and 1% glutamax (Gibco, #35050-061) until sufficiently expanded. Mouse CCL206 lung fibroblasts (CCL-206; ATCC, Wesel, Germany, RRID: CVCL_0437) were cultured in DMEM (Lonza, Basel, Switzerland, #42430):Ham’s F12 (Gibco, #21765) (1:1) supplemented with 10% FBS, 2 mM L-glutamine (Life Technologies under Thermo Fisher Scientific, Waltham, MA, USA, #35050–061), 100 U/ml penicillin/streptomycin, and 1% amphotericin B (Gibco, #15290-026). All cells were maintained at 37°C with 5% CO_2_ and confirmed to be mycoplasma negative when used.

### Mouse epithelial cell isolation

Epithelial (CD31^−^/CD45^−^/EpCam^+^) cells were isolated as previously described [25]. C57BL/6J mice were anesthetized by injecting a mixture of 400 mg/kg Ketamidor^®^ and 1 mg/kg Dexdomitor^®^ subcutaneously. After confirming successful anesthesia, the animal was euthanized by exsanguination, after which the lungs were flushed through the heart with PBS. Subsequently, the lungs were filled with 1.5 ml dispase (BD Biosciences, Oxford, UK, #354235), then removed from the thoracic cavity, and placed into an additional 1 ml of dispase. They were incubated at room temperature for 45 minutes. Next, the trachea was removed, and lung lobes were homogenized in DMEM with DNase1 (Applichem, Darmstadt, Germany #A3778). The resulting suspension was filtered using cell strainer filters (Corning Incorporated, Corning, NY, USA, #431752), incubated with microbeads conjugated to antibodies for CD45 (Miltenyi Biotec, Teterow, Germany #130-052-301, RRID: AB_2877061) and CD31 (Miltenyi, #130-097-418, RRID: AB_2814657), and loaded on LS columns (Miltenyi #130-091-051). Positive selection for epithelial cells was performed on the CD31^−^/CD45^−^ suspension using EpCam (CD326) microbeads (Miltenyi #130-105-958). EpCam^+^ cells were resuspended in DMEM :100:342 100:342 100:342 Ham’s F-12 (1:1) with 10% FBS, 2 mM L-glutamine, 100 U/ml penicillin/streptomycin, and 1% amphotericin B.

### Organoid culture

The organoid assay was based on published protocols [25]. In this manuscript, 2 types of organoids were used; mouse organoids, composed of primary mouse EpCam^+^ cells combined with CCL206 fibroblasts, and mixed organoids formed by primary mouse EpCam^+^ cells co-cultured with primary human fibroblasts. Primary human lung fibroblasts of passage 5 until 7 (table 2) or CCL206 fibroblasts were proliferation-inactivated with mitomycin C (10 μg/ml, Sigma-Aldrich, St. Louis, MO, USA, #M4287) for 2 hours, after which they recovered in normal medium for at least 1 hour, and were subsequently trypsinized. 10,000 fibroblasts were mixed with 10,000 freshly isolated mouse lung EpCam^+^ cells. They were seeded into transwell inserts for 24-well plates (Thermo Fischer Scientific, Waltham, USA #10421761) in 100 μl growth factor-reduced Matrigel (Fisher Scientific, Landsmeer, The Netherlands #11523550) diluted 1:1 with DMEM:Ham’s F12 (1:1) containing 10% FBS, 2 mM L-glutamine, 100 U/ml penicillin/streptomycin, and 1% amphotericin B. Cultures were maintained in DMEM/F12 with 5% FBS, 2 mM L-glutamine, antibiotics, insulin-transferrin-selenium (Gibco #15290018), recombinant mouse EGF (0.025μg/ml, Sigma-Aldrich, #SRP3196) and bovine pituitary extract (30μg/ml, Sigma-Aldrich, #P1476) at 37°C with 5% CO_2_. Y-27632 (10 μM, Tocris Bioscience, Oxford, UK, #1254) was added for the first 48 hours of culture. Media were refreshed every 2-3 days. Both mouse and mixed organoids were exposed to recombinant human IL-11 (rhIL-11, R&D systems, Minneapolis, MN, USA, #218-IL) during the entire organoid culture period of 14 days.

**Table 2:** patient characteristics of primary human fibroblasts used for organoid culture

| **Patient group** | **Sex M/F** | **Age Median (range)** | **Smoking status Never/ex/current** | **Pack years Median (range)** | **FEV1 (% of predicted pre BD) Median (range)** | **FVC (% of predicted pre BD) Median (range)** |
| --- | --- | --- | --- | --- | --- | --- |
| Control | 3/2 | 49.5 (46-69) | 1/2/1 | Ex: 26 (20-32)<br>Current: 33 | 96.9 (90.7-105) | 104.5 (104-105)<br>3 unknown |
| IPF | 4/1 | 63 (36-68) | 1/3/0<br>1 unknown | Ex: 20 (15-25)<br>3 unknown | 61 (45-81)<br>1 unknown | 51 (43-83)<br>1 unknown |

### Organoid immunofluorescence

Organoid cultures were fixed with ice-cold acetone/methanol (1:1) for 12 minutes at −20 °C and stored in 0.02% sodium azide in PBS at 4 °C until further use. Vehicle control and treatment groups were collected and stained simultaneously. Cultures were blocked with 5% BSA for 2 hours at room temperature, and subsequently incubated with primary antibodies rabbit anti-Prosurfactant Protein C (pro-SPC) (MilliporeSigma, Burlington, MA, USA, #AB3786, RRID: AB_91588) and mouse anti-Acetylated α Tubulin (ACT) (Santa Cruz Biotechnology, Dallas, TX, USA, #sc-23950, RRID: AB_628409) diluted 1:200 in 0.1% BSA 0.1% Triton-X100 in PBS at 4 °C overnight. They were then washed in PBS, and incubated with secondary antibodies donkey anti-rabbit Alexa fluor 488 (Thermo Fisher Scientific #A21206, RRID: AB_2535792) and donkey anti-mouse Alexa fluor 568 (Thermo Fisher Scientific #A10037, RRID: AB_2534013) for 2 hours at room temperature. Cultures were washed, excised and mounted on glass slides with mounting medium containing 4⍰,6-diamidino-2-phenylindole (DAPI) (Abcam, #104139). Descriptive immunofluorescence images were taken using a Leica SP8 confocal microscope at magnification 20x or 40x. For quantification, one individual scored the counts who was blinded to the samples. At least 50 organoids per culture were assessed using a Leica DM4000B fluorescence microscope at magnification 40x.

### Precision cut lung slices

C57BL/6J mice were anesthetized by subcutaneous injection of 400 mg/kg Ketamidor^®^ and 1 mg/kg Dexdomitor®. After anesthesia was confirmed, the animal was euthanized by exsanguination, after which the lungs were filled until just inflated with 1.5% low-melting point agarose (Gerbu Biotechnik GmbH, Wieblingen, Germany) solution in CaCl_2_ (0.9 mM), MgSO_4_ (0.4 mM), KCl (2.7 mM), NaCl (58.2 mM), NaH_2_PO_4_ (0.6 mM), glucose (8.4 mM), NaHCO_3_ (13 mM), HEPES (12.6 mM), sodium pyruvate (0.5 mM), glutamine (1 mM), MEM-amino acids mixture (1:50) and MEM-vitamins mixture (1:100) pH=7.2. After inflation, lungs were removed and placed on ice for 15 minutes to allow the agarose to solidify. Subsequently, the lungs were separated in lobes and placed into cores. A tissue slicer (CompresstomeTM VF-300 microtome, Precisionary Instruments, San Jose, CA, USA) was used to cut precision cut lung slices (PCLS) of 250 μm in thickness in CaCl_2_ (1.8 mM), MgSO_4_ (0.8 mM), KCl (5.4 mM), NaCl (116.4 mM), NaH_2_PO_4_ (1.2 mM), glucose (16.7 mM), NaHCO_3_ (26.1 mM), HEPES (25.2 mM), pH=7.2. Slices were washed 4 times for 30 minutes (2 hours total). Three slices were placed in one well of a 12-well plate and incubated with 2 mL DMEM (Lonza, #42430) with 0.6% amphotericin B, 1 mM sodium pyruvate (HyClone Laboratories, Logan, UT, USA, #SH30239.01), 1% MEM non-essential amino acids (Gibco, #11130036), 0.6% gentamycin (Sigma-Aldrich, #G1272) and 100 U/ml penicillin/streptomycin, and 100 ng/ml rhIL-11 for 48 hours. For gene expression studies, the slices were collected and stored at −80 °C until use.

### Gene expression analyses

For gene expression studies, 300,000 primary fibroblasts of control and IPF donors (table 3) of passage 5 until 7 were seeded in 6-well culture plates with low glucose DMEM (Biowest, Nuaillé, France, #L0064-500), supplemented with 10% FBS, 100 U/ml penicillin/streptomycin and 1% glutamax. The cells were allowed to settle for 24 hours and were subsequently serum deprived for 24 hours with low glucose DMEM with 0.1% BSA, glutamax and antibiotics. Cells were then exposed to rhIL-11 (R&D systems, #218-IL) for 24 hours, after which they were collected in TRIzol (Invitrogen under Thermo Fisher Scientific, Waltham, MA, USA, #15596018). Total RNA was extracted from primary human lung fibroblasts according to manufacturer’s instructions. The Maxwell simplyRNA tissue kit (Promega, Madison, WI, USA, #AS1280) and Maxwell 16 Instrument (Promega) were used to isolate RNA from mouse PCLS according to manufacturer’s instructions. Total RNA concentrations were determined with a NanoDrop ND-1000 spectrophotometer. Equal amounts of total mRNA were then reverse transcribed (Promega). Real time PCR was performed with SYBR green as the DNA binding dye (Roche Applied Science, Mannheim, Germany) on a 7900HT Fast Real-Time PCR System (Applied Biosystems under Thermo Fisher Scientific, Waltham, MA, USA), with denaturation at 94°C for 30 seconds, annealing at 60°C for 30 seconds and extension at 72°C for 30 seconds for 40 cycles followed by 5 minutes at 72°C. Gene expression was normalized to *B2M*, *SDHA* and *HMBS* for human samples (primary fibroblasts) and to *Rpl13a*, *B2m* and *Actb* for mouse samples (PCLS). Fold changes were calculated using the 2^−ΔΔCT^ method. The primers used are listed in tables S1 and S2.

**Table 3:**
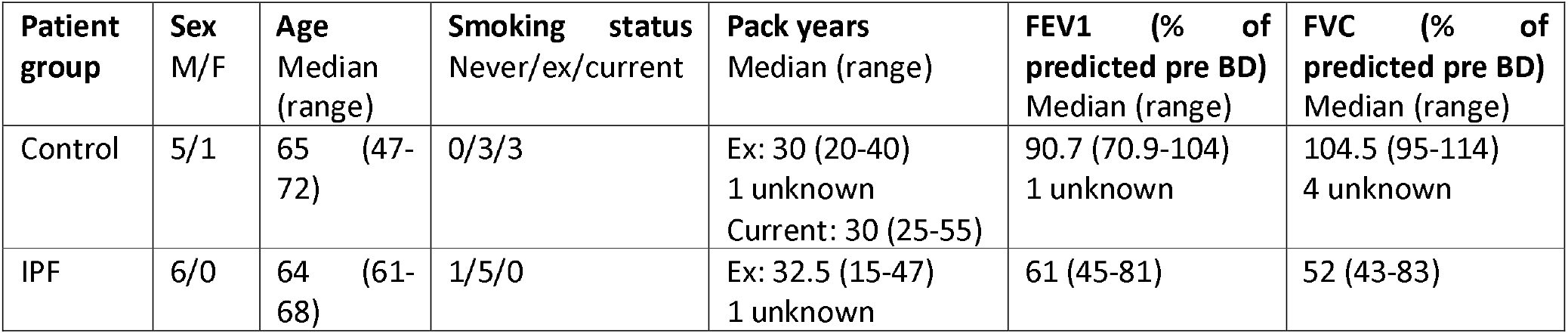
patient characteristics of primary human fibroblasts used for gene expression studies

### RNA-sequencing

Primary human lung fibroblasts of control donors (see table 4) of passage 5 until 7 were seeded at a density of 300,000 cells per well in 6-well culture plates, allowed to settle for 24 hours, serum-starved for 24 hours and then exposed to 100 ng/ml rhIL-11 for 24 hours. The fibroblasts were then lysed in 1 mL of TRIzol reagent and total RNA was isolated according to the manufacturer’s instructions. Total RNA concentrations were initially determined with a NanoDrop ND-1000 spectrophotometer and quantified in detail using Bio-analyzer fragment analyser from Agilent. An Illumina NovaSeq 6000 sequencer was used for bulk RNA-seq data analysis by GenomeScan (the Netherlands). The procedure included data quality control, adapter trimming, alignment of short reads and feature counting. Library preparation was checked by calculating ribosomal (and globin) content. Checks for possible sample and barcode contamination were performed and a set of standard quality metrics for the raw data set was determined using quality control tools (FstQC v0.34 and FastQA). Prior to alignment, the reads were trimmed for adapter sequences using Trimmomatic v0.30. To align the reads of each sample, the human reference Ensembl GRCh37.75 was used. Gene duplicates were removed. Counts were normalized in R 4.1.0 and analyzed using paired-sample analysis in the DESeq2 package 1.34.0 in R to determine differentially expressed genes (DEGs), which were considered significant when padj<0.05. Due to the low number of statistically significant DEGs, all significant and non-significant DEGs were used for pathway enrichment analysis using the fgsea package 1.20.0 in R, and pathways were considered significant when padj<0.05. A volcano plot was generated using the Enhanced Volcano package in R, with a cut-off value of 0.5 for Log2Fold change and 0.05 for padj value. The complete dataset is available as a supplementary table via www.figshare.com/s/59954d2189db9288be7b.

**Table 4:**
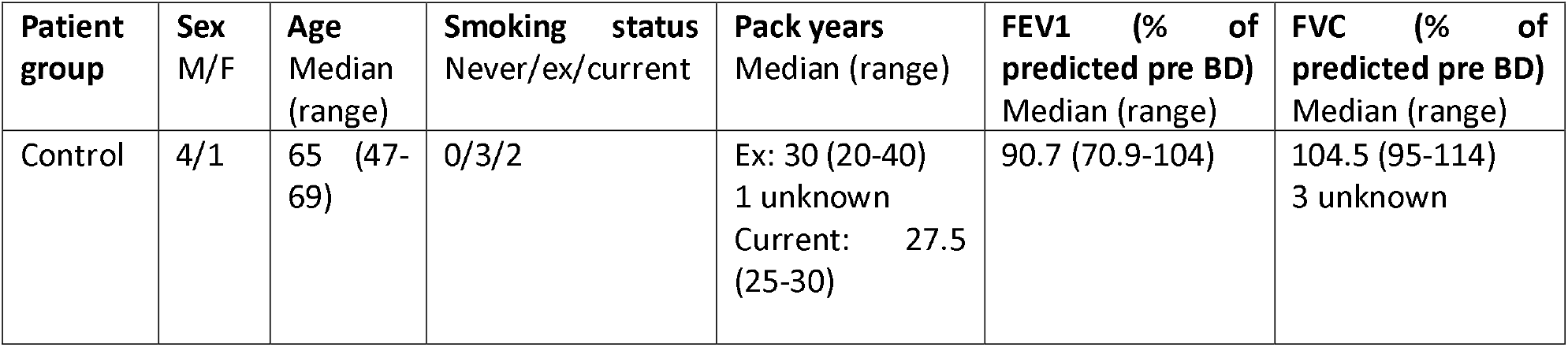
patient characteristics of primary human fibroblasts used for RNA-sequencing

| Patient group | Sex M/F | Age Median (range) | Smoking status Never/ex/current | Pack years Median (range) | FEV1 (% of predicted pre BD) Median (range) | FVC (% of predicted pre BD) Median (range) |
| --- | --- | --- | --- | --- | --- | --- |
| Control | 4/1 | 65 (47-69) | 0/3/2 | Ex: 30 (20-40)<br>1 unknown<br>Current: 27.5 (25-30) | 90.7 (70.9-104) | 104.5 (95-114)<br>3 unknown |

### Data analyses

For sequencing data, statistical analyses were performed in R as described above. Statistical evaluation of all other data were performed using GraphPad Prism 8. Data are presented with mean and standard deviation unless described otherwise in figure legends, and N refers to the number of biological replicates (number of animals or individual donors). Prior to the studies, power calculations were performed using organoid number as primary read-out parameter at α 0.05 and power 0.8. A total of 10 biological replicates were found to be necessary. No exclusion criteria were set, and as such no animals or data points were excluded from any studies described here. Data were checked for normality using the Shapiro-Wilk test. Data that failed the normality test were log transformed and tested again. Data that were normalized to vehicle control were always first log transformed before testing for normality and performing statistical tests. For normally distributed data, a two-tailed t-test was used to compare 2 groups, whereas a one-way ANOVA was used to compare more than 2 groups, and a two-way ANOVA was performed when data were split into subgroups. For not normally distributed data with 2 groups, a Wilcoxon test was used, and more than 2 groups were compared using a Kruskall-Wallis test. The statistical tests used are also specified in the figure legends. Differences were considered statistically significant when p<0.05.

## Results

### IL-11 and IL-11R expression in human lung tissue

The cell type specific expression of IL-11 and IL-11 receptor (IL-11R) in the human lung remains incompletely understood, though publicly available RNA sequencing datasets suggest fibroblasts, myofibroblasts, innate lymphoid cells, endothelial cells and various epithelial cells express IL-11, whereas IL-11R appears to be expressed by a wide variety of cells, including numerous epithelial cell types, fibroblasts, smooth muscle cells, dendritic cells and endothelial cells (figure S1). We aimed to further investigate the expression pattern on protein level by performing immunohistochemical staining for IL-11 and IL-11R in human lung tissue. First, secondary antibody specificity was confirmed by excluding the IL-11 primary antibody from the staining procedure (figure 1A). IL-11 protein expression was observed in the airway epithelium, a subset of macrophages and to a lesser extent in the tunica media of blood vessels and in the smooth muscle around airways (figure 1B-D). In IPF tissue, there was also some IL-11 staining in AT2 cells in areas of AT2 cell hyperplasia (figure 1E). IL-11 protein expression was not found in fibroblast foci in IPF tissue. Quantitative analysis of IL-11 tissue positive area (figure 1F) and average intensity of staining (figure 1G) indicated that there were no overall differences between control and IPF tissue (p=0.2401 and p=0.8256 respectively). Secondary antibody specificity was also validated for IL-11R staining (figure 1H). IL-11R staining was primarily visible in macrophages and in smooth muscle surrounding vessels and airways, and in IPF tissue, also in randomly organized smooth muscle. IL-11R was also present to a lesser extent in airway epithelium and the endothelium (figure 1I-L). IL-11R staining was not observed in fibroblast foci in IPF tissue. The IL-11R staining area was significantly decreased in IPF lung tissue compared with control tissue (figure 1M) (p=0.0209), whereas the intensity of IL-11R staining did not differ between groups (p=0.7016) (figure 1N). For the quantification of the immunohistochemical staining of both IL-11 and IL-11R, the variability within each group was substantial. Indeed, numerous control and IPF tissue sections were almost completely negative for IL-11 and/or IL-11R staining, whereas others did show positivity. Interestingly, we noticed heterogeneity not only within the groups, but even within the same tissue, as structures were mostly only partially positive for IL-11 and IL-11R (figure S2-3). For example, generally only a subset of macrophages and AT2 cells were positive within one tissue section, and protein expression could be found in the epithelium of one airway, whereas other airways present in the tissue were negative. Due to the inconsistent protein expression pattern of both IL-11 and IL-11R, no differences in expression pattern were found between control and IPF tissue.

**Figure 1:**
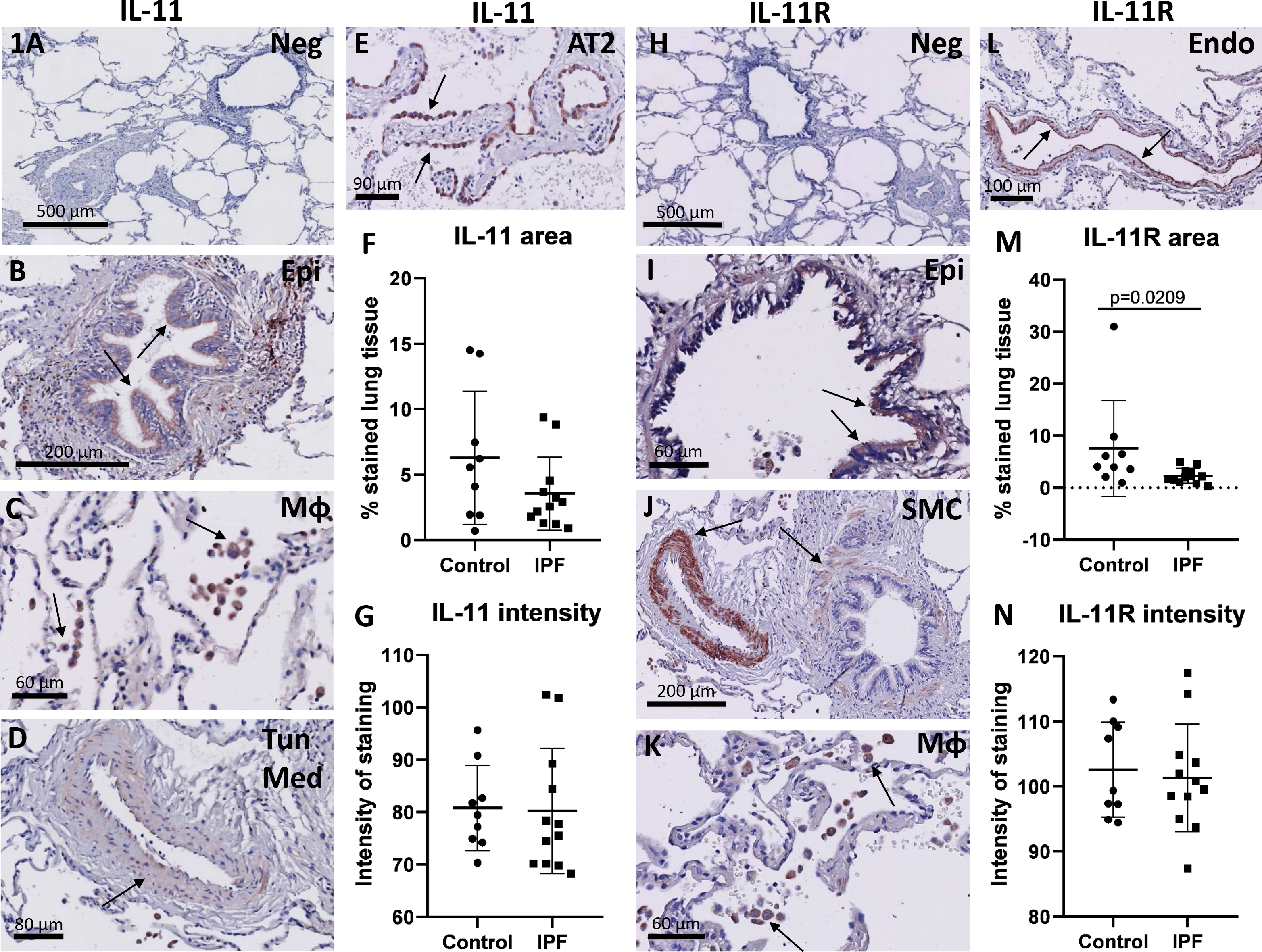
expression pattern of IL-11 and IL-11R in the human lung. Human lung tissue sections of control and idiopathic pulmonary fibrosis (IPF) donors were immunohistochemically stained for IL-11 and IL-11R, developed with NovaRED (red) and counterstained with hematoxylin (blue). Arrows indicate positive cells or structures. Negative control image of IL-11 staining (A) was obtained by omitting primary antibody from immunohistochemistry procedure. IL-11 staining was present in airway epithelium (B), macrophages (C) and the tunica media of blood vessels (D). Example images B-D are from the control group. In IPF, IL-11 staining was also present in areas of alveolar type 2 (AT2) cell hyperplasia (E). Analysis of IL-11 staining area (F) and intensity (G) in control (N=9) and IPF (N=12) tissue, unpaired t-test on log transformed data. Negative control image of IL-11 receptor (IL-11R) staining (H). IL-11R expression was observed in airway epithelium (I), smooth muscle (J), macrophages (K) and the endothelium (L). Example images I-L are from the control group. Analysis of IL-11R staining area (M) and intensity (N) in control (N=9) and IPF (N=12) tissue, unpaired t-test on log transformed data.

### IL-11 negatively impacts progenitor activation and alveolar differentiation

Since little is known on the influence of IL-11 on alveolar epithelial progenitor cell function, we used a mouse organoid model composed of primary mouse EpCam^+^ cells combined with CCL206 mouse lung fibroblasts (figure 2A) to study the regenerative capacity of epithelial progenitor cells in response to IL-11 (figure 2B). Overall, the number of organoids formed on day 14 was significantly decreased in the presence of rhIL-11 (p=0.0045). Post-hoc analyses showed that 10 ng/ml and 100 ng/ml IL-11 both induced a significant reduction in the number of organoids formed (p=0.0303 and p=0.0224 respectively) (figure 2C). The organoid number data demonstrated remarkable variability, indicating a diversity of response with several organoid cultures not, or only minimally inhibited by IL-11 and others that were extensively inhibited. We hypothesized this may be caused by the sex of the mice from which the epithelial cells were isolated. We then expanded the organoid experiments and stratified the organoid number results for mouse sex from which the epithelial cells were isolated (figure 2D). Although IL-11 significantly reduced the number of organoids formed (p=0.0017), the response to IL-11 was not dependent on the sex of the donor mouse.

**Figure 2:**
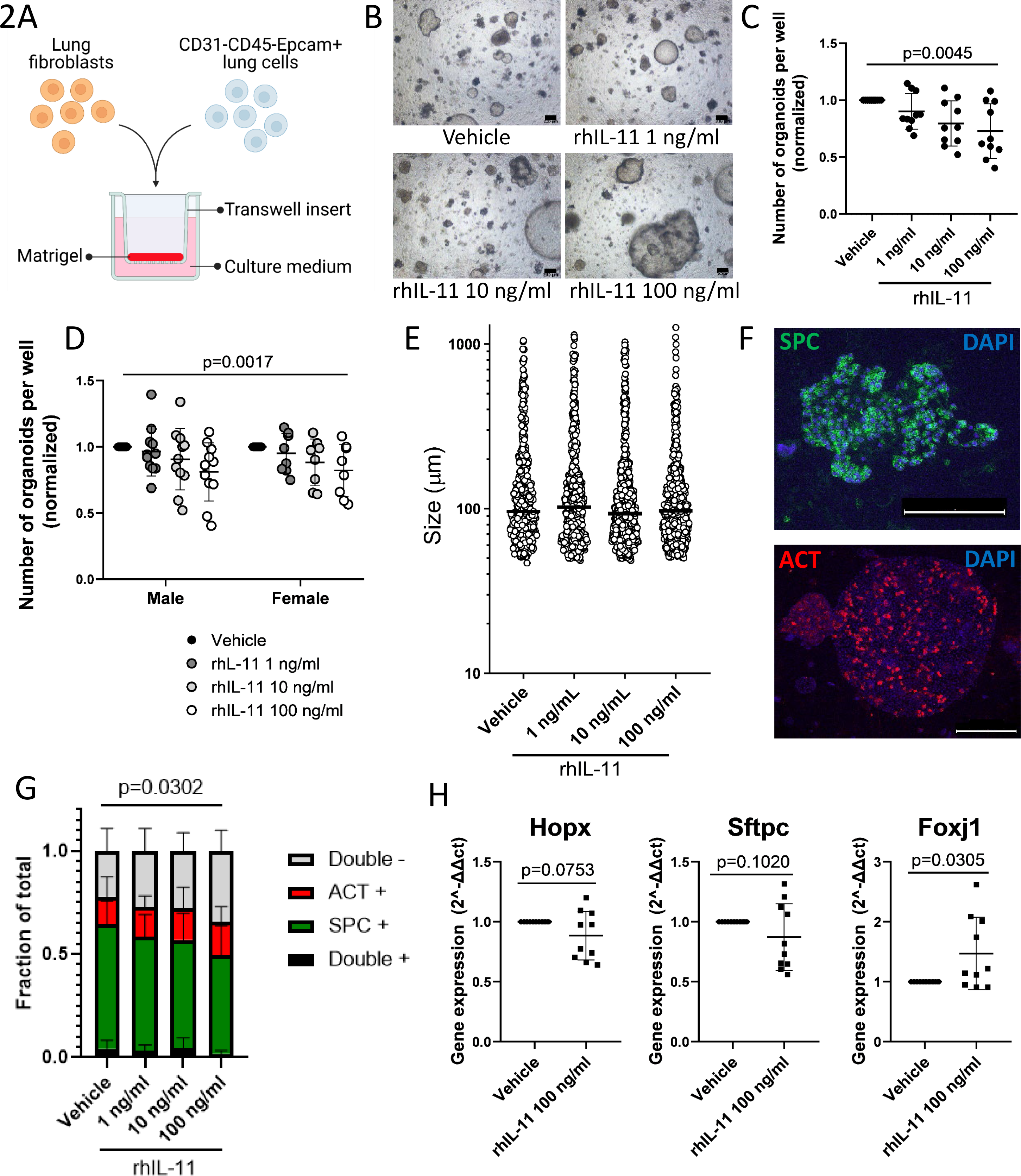
rhIL-11 dysregulates epithelial progenitor cell function in mouse organoids. (A) Primary mouse EpCam cells were combined with CCL206 mouse lung fibroblasts, seeded in matrigel and allowed to form organoids during 14 days in the presence of rhIL-11 (created with Biorender.com). (B) Representative Brightfield images of mouse organoids exposed to rhIL-11. Scale bar = 200 μm. (C) Normalized mouse organoid number on day 14 after continuous treatment with a dose curve of rhIL-11 (N=10, One-way ANOVA with Sidak’s post hoc test on log transformed data). (D) Normalized number of mouse organoids in response to rhIL-11, separated by mouse sex (N=11 male, N=8 female, Two-way ANOVA with Sidak’s post hoc test on log transformed data). (E) Organoid diameter on day 14 in response to rhIL-11, median is shown (N=10, Kruskall-Wallis test with Dunn’s post hoc test). (F) Representative fluorescent images of a Prosurfactant protein C (pro-SPC)+ (scale bar = 50 μm) and Acetylated α Tubulin (ACT)+ (scale bar = 100 μm) organoid. (G) Quantification of the fraction of pro-SPC-expressing and ACT-expressing mouse organoids in response to rhIL-11 (N=10, Two-way ANOVA with Sidak’s post hoc test). (H) Precision cut lung slices of wild type mice were exposed to 100 ng/ml rhIL-11 for 48 hours, after which RNA was isolated from the whole slice and gene expression studies were performed. Gene expression of alveolar epithelial cell markers Hopx and Sftpc, and of ciliated cell marker Foxj1, N=10, paired t-test on log transformed data.

The median organoid diameter on day 14 was not altered in the presence of IL-11 (p=0.3819) (figure 2E). Immunofluorescence was performed for differentiation markers pro-SPC and ACT, marking AT2 cells and ciliated cells respectively (figure 2F). Quantification of the organoid expression of pro-SPC and ACT revealed that IL-11 significantly influenced marker expression (p=0.0302). 100 ng/ml rhIL-11 induced a reduction of the pro-SPC expressing organoid fraction (p=0.0089) and concurrently an increase in the double negative fraction (pro-SPC-ACT-organoids) (p=0.0126) (figure 2G).

After observing indications of progenitor cell dysfunction induced by IL-11 in the organoid model, we assessed whether IL-11 could also affect the epithelium in the model of mouse precision cut lung slices (PCLS), in which most lung cell types are present in their physiological environment, and natural cell-cell and cell-extracellular matrix interactions are preserved. After incubating the slices with rhIL-11 for 48 hours, we assessed the gene expression of epithelial cell markers. Interestingly, the expression of alveolar cell markers (*Hopx* and *Sftpc*) was non-significantly downregulated by IL-11 (p=0.0753 and p=0.1020 respectively) (figure 2H), which is in line with the mouse organoid data for which IL-11 induced a decrease in pro-SPC protein expression. Whereas the expression of *Sgb1a1* and *Muc5ac* was unaffected by IL-11, *Foxj1* expression was significantly increased (p=0.0305) (figure S4A, figure 2H). To examine any pro-fibrotic effects of IL-11, we assessed expression of genes associated with fibrosis (figure S4B). Most fibrosis marker genes were not altered in response to IL-11, though *Tgfb1* expression was downregulated (p=0.0039). The expression of both *Il11* and *Il11ra1* in response to IL-11 stimulation was unchanged (figure S4C). Similar to the organoid number data described in figure 2C, the expression of various marker genes in the PCLS studies showed large variability, in which generally marker expression in PCLS originating from several mice was considerably affected by IL-11 exposure whereas PCLS of other mice gave only a minimal response.

Next, we wondered if the IL-11 induced aberrant progenitor response that we observed in mouse organoids would also occur using a combination of primary mouse EpCam+ cells and primary human fibroblasts (figure 3A), and whether the disease state of the fibroblasts would affect the response to IL-11. Importantly, the co-culture of primary mouse EpCam+ cells and primary human fibroblasts was previously shown to yield organoids comparable in number, size and differentiation to those formed by primary mouse EpCam+ cells combined with CCL206 fibroblasts [25]. Overall, rhIL-11 exposure induced a significant decrease in the number of organoids (p=0.0001) formed by primary mouse EpCam+ cells combined with primary human fibroblasts (figure 3B). This was not dependent on whether the fibroblasts used in the cultures were isolated from control or IPF tissue (p=0.3055), indicating the disease state of the fibroblasts does not affect the response to IL-11 in the organoid culture. There was no effect on organoid diameter in cultures established with either control or IPF fibroblasts (figure 3C-D), similar to our previous results.

**Figure 3:**
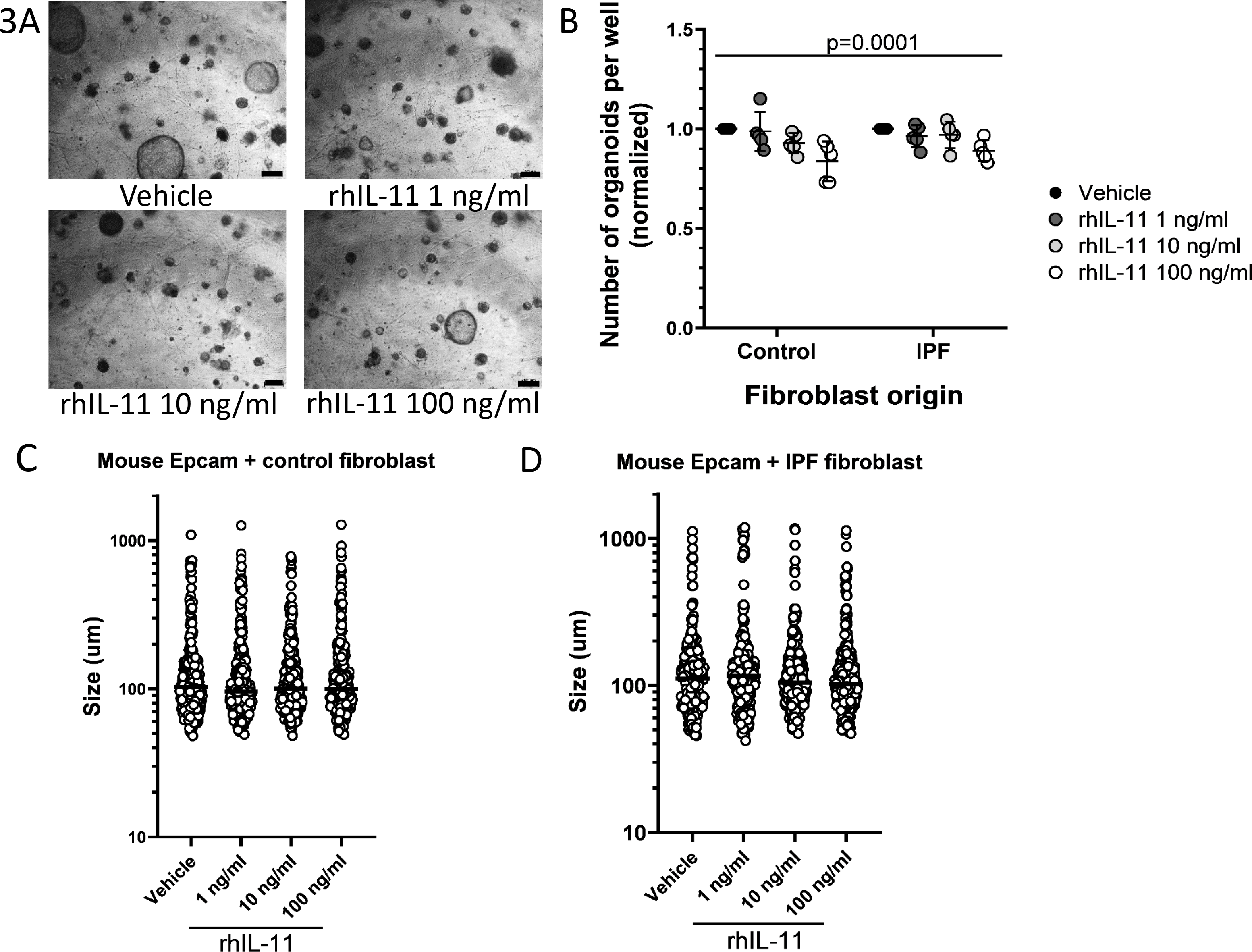
the mouse epithelial progenitor cell response is disturbed by IL-11 in mouse/human organoid co-cultures. Primary mouse EpCam cells mixed with primary human lung fibroblasts (either control or idiopathic pulmonary fibrosis (IPF)) were seeded in matrigel, cultured for 14 days and continuously exposed to rhIL-11; resulting organoid formation was studied. (A) Representative Brightfield images of organoids formed by mouse EpCam+ cells with control primary human fibroblasts, in response to rhIL-11. Scale bar = 250 μm. (B) Normalized number of mouse/human organoids in response to 14 day treatment with rhIL-11 (N=5 control fibroblasts, N=5 IPF fibroblasts, Two-way ANOVA with Sidak’s post hoc test on log transformed data). (C) Diameter of organoids formed by mouse EpCam+ cells with control fibroblasts after rhIL-11 treatment, median is shown (N=5, Kruskal-Wallis test with Dunn’s post hoc test). (D) The effect of rhIL-11 on the diameter of organoids formed by mouse EpCam+ cells with IPF fibroblasts, median is shown (N=5, Kruskal-Wallis test with Dunn’s post hoc test).

### The production of known epithelial progenitor supportive factors by fibroblasts is not affected by IL-11

Since the organoid culture is a co-culture system, the effect of IL-11 on alveolar progenitors (figure 2 and 3) could be indirect and regulated by a fibroblast response. To assess their response to IL-11, we exposed CCL206 mouse lung fibroblasts to both recombinant mouse IL-11 (rmIL-11) and rhIL-11 for 30 minutes, after which cells were lysed and subjected to western blot. An increase in phosphorylated STAT3 and ERK1/2 was observed in CCL206 fibroblasts after rmIL-11 and rhIL-11 treatment, where the response to rhIL-11 was more pronounced, indicating activation of downstream signaling pathways JAK/STAT3 and MEK/ERK (figure S5A-C). The activation of downstream signaling, particularly MEK/ERK, by IL-11 in primary human fibroblasts has been described previously [12]. Next, we studied whether IL-11 could affect the production of progenitor supportive factors by fibroblasts. Primary human fibroblasts (either control or IPF) were exposed to IL-11 for 24 hours after which gene expression studies were performed. The expression of progenitor supporting factors in response to rhIL-11 (figure S6A) was variable, with no significant effects on any measured genes encoding secreted factors. We also determined the expression of the fibrosis-marker gene *Fibronectin 1 (FN1)* (figure S6B), which was not significantly affected by IL-11. Since little is known about the regulation of IL-11 and IL-11R expression [26], the effect of IL-11 itself on this expression was also assessed. In addition, TGFβ is an important inducer of IL-11, and an IL-11 antibody was found to inhibit the phosphorylation of SMAD2 in a bleomycin mouse model, indicating modulation of TGFβ signaling by IL-11 [12]. Thus, we asked whether IL-11 could modulate TGFβ by affecting its gene expression. The expression of *TGFB1*, *IL11* and *IL11RA* was, however, not influenced by IL-11 treatment of either control or IPF fibroblasts (figure S6B-C).

### IL-11 affects processes in fibroblasts including metabolism, cellular stress and cross-talk mechanisms

As we did not find IL-11-induced gene expression changes of known epithelial cell supportive factors, we performed bulk RNA-sequencing in order to get a broader overview of the influence of IL-11 on primary human lung fibroblasts. The volcano plot (figure 4A) shows there were relatively few genes that were differentially expressed in response to 24 hour exposure to rhIL-11 (24 genes with padj<0.05), indicating IL-11 may not induce a response as strong as for example TGFβ (previously found to induce 3795 differentially expressed genes in primary human fibroblasts with FDR<0.01) [25]. Significantly upregulated genes include *Hexokinase 2 (HK2)* and *Lactate Dehydrogenase A (LDHA)* which are involved with glycolysis, and *Suppressor Of Cytokine Signaling 3 (SOCS3)* which is a target gene of STAT3 and acts as a negative feedback mechanism of JAK/STAT signaling [27]. The gene expression of *Elastin (ELN)*, a ECM component which provides elasticity, is significantly downregulated in IL-11 exposed fibroblasts (figure 4A). Due to the limited number of differentially expressed genes, we performed Gene Set Enrichment Analysis (GSEA) on all genes (including not significant genes) to increase the power. This analysis revealed IL-11 significantly upregulated several pathways (figure 4B), including multiple pathways associated with the regulation of proliferation (*E2 factor (E2F) targets*, *G2M checkpoint*, *MYC targets* and *mitotic spindle*), which was previously shown to be increased in fibroblasts by IL-11 [28], and downstream mechanisms of IL-11 signaling; JAK/STAT3, MEK/ERK (KRAS) and PI3K/Akt/mTOR [27], indicating IL-11 does indeed induce expected responses in primary fibroblasts. Other processes that were affected by IL-11 include *TGFβ signaling*, inflammation (*TNFα signaling via NF-κB*, *inflammatory response*, *IFNγ response*, *complement* and *IFNα response*), cellular stress (*unfolded protein response* and *reactive oxygen species pathway*), metabolism (*glycolysis*) and cross-talk pathways (*Notch signaling*).

**Figure 4:**
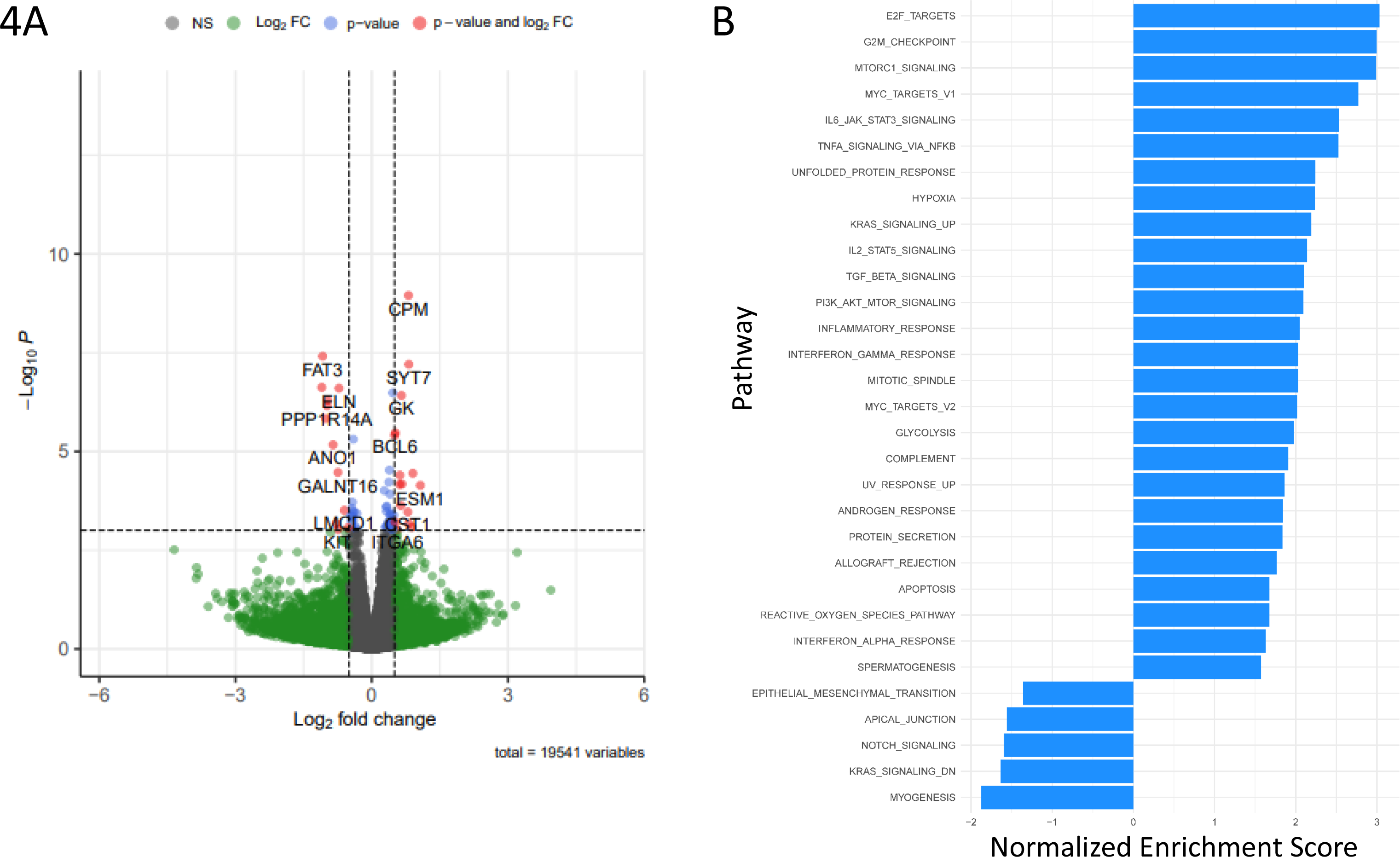
IL-11 alters several mechanisms in primary human fibroblasts which can influence fibrosis and cellular cross-talk. Primary human fibroblasts of control donors (N=5) were treated with 100 ng/ml rhIL-11 for 24 hours, and subsequently subjected to bulk RNA-sequencing analysis. (A) Volcano plot showing significantly altered genes in control fibroblasts treated with IL-11 (Log2fold change cut-off = 0.5 and padj cut-off = 0.05). (B) Gene set enrichment analysis (GSEA) showing upregulated pathways (positive enrichment score) and downregulated pathways (negative enrichment score) induced by IL-11.

## Discussion

In this study we describe for the first time the distribution of IL-11 and IL-11R in human lung tissues on protein level, and show that IL-11 plays important roles in lung epithelial cell growth and differentiation. IL-11 immunostaining is present in various cell types and structures, including macrophages and AT2 cells, and IL-11R staining was mostly present in smooth muscle and macrophages, but also in other areas. Furthermore, we showed that IL-11 affects epithelial progenitor cell function by inhibiting progenitor activation and inducing an imbalance of epithelial differentiation. Surprisingly, IL-11 did not induce changes in fibrotic marker gene expression in primary human lung fibroblasts or PCLS. Finally, we explored various mechanisms by which fibroblast epithelial cross-talk may be disturbed by IL-11, possibly contributing to defective alveolar repair. Previous studies indicated IL-11 receptor expression is particularly high in stromal cells, including fibroblasts and smooth muscle cells (SMCs), though epithelial and immune cell lineages also express IL-11R [7,9,29–31]. Data in the IPF cell atlas suggest basal gene expression of IL-11R in the human lung is limited, but fibroblasts, smooth muscle cells, endothelial cells, immune cells and various epithelial cells appear to express IL-11R to a higher extent than other cells [32,33]. Here we observed immunohistochemical staining of the receptor primarily in smooth muscle and macrophages, and to a lesser extent in the airway epithelium and the endothelium, but not in fibroblast foci in IPF tissue. The discrepancy between literature and this study might be caused by previous studies mainly reporting gene expression data, often in cell cultures or mouse organs, whereas here protein expression in human lung tissue was assessed. Quantitative analysis of IL-11R staining indicated a significant decrease in percentage of tissue area positive for IL-11R in IPF compared to control. The reason for this decrease is not known since the regulation of the expression of IL-11R is largely unexplored [26], though the disorganization of the lung architecture with a different composition of cells and ECM in IPF compared to healthy lung tissue may affect the overall expression of IL-11R as well. Though IL-11R expression was not observed in fibroblast foci, we and others have shown that fibroblasts activate downstream signaling pathways in response to IL-11, including MEK/ERK and JAK/STAT [12]. This suggests IL-11R expression is dynamically regulated and may differ between cellular subsets or activation states, highlighting the need for studies on IL-11R regulation. Whereas IL-11 mRNA overexpression has been reported in IPF lungs, and a small study (N=3) also showed an increase in immunohistochemical staining for IL-11 in IPF [12], here we found no differences between the percentage area nor intensity of IL-11 staining when comparing control and IPF tissues. In both previous studies and this study, the data were not corrected for smoking status. Since the influence of smoking on the expression of IL-11 is of yet unknown, the smoking status of the IPF and control groups may affect observed differences in IL-11 expression. Additionally, whether the IPF lung samples contained active or only established fibrosis could influence the expression of IL-11 in this study and results described in literature. Lung fibroblasts, SMCs, epithelial cells and eosinophils were previously reported to produce IL-11 *in vitro* [34–37], whereas damaged epithelial cells and fibroblasts have been suggested as primary sources of IL-11 in the human lung *in vivo* [12,16,30,38]. Here, we observed IL-11 in the airway epithelium, in AT2 cells in areas of AT2 cell hyperplasia, and in SMCs. Interestingly, IL-11 expression was also present in macrophages. Recent studies suggest macrophages are of importance in the pathogenesis of IPF, as they were proposed to contribute to fibrosis and dysfunctional epithelial repair [39–41]. We also noted expression of IL-11 in AT2 cells in areas of AT2 cell hyperplasia (figure 1E), but AT2 cells in control lung were generally negative. This is of particular interest since epithelial cells were previously proposed to produce IL-11 [12,16] and epithelial cell dysregulation relates to the initiation of fibrosis [16].

Next, we showed that IL-11 negatively affected organoid formation. The effect of IL-11 on lung epithelial cells has been of limited research focus, though it was recently shown that both A549 cells and H1792 cells are able to respond to IL-11 by activating downstream ERK signaling [42]. Additionally, IL-11 has been studied in human embryonic stem cell (hESC) derived organoids that model Hermansky-Pudlak syndrome associated interstitial pneumonia (HPSIP), which also recapitulate features of IPF. These organoids had an abnormal morphology, and overexpressed IL-11 specifically in the epithelial fraction. Interestingly, IL-11 knockout HPSIP organoids had a normal morphology corresponding to wild type organoids, suggesting IL-11 negatively impacts epithelial cells [16]. Our organoid data reveal that IL-11 decreases the number of organoids formed, indicating disrupted progenitor cell activation. IL-11 also reduces the pro-SPC+ organoid fraction, which most likely indicates dysfunction of AT2 cells or disturbances in the differentiation towards alveolar cell types. Primary mouse AT2 cells treated with IL-11 were recently reported to have an increased expression of KRT8 and stalled transdifferentiation into AT1 cells. IL-11 was also shown to inhibit primary human alveolar cell proliferation. Moreover, the accumulation of KRT8+ cells and delayed AT2-AT1 cell transdifferentiation after bleomycin injury was rescued by specific Il11ra1 deletion in AT2 cells, supporting our hypothesis that IL-11 induces AT2 cell dysfunction and impaired alveolar differentiation [43]. Importantly, dysregulated AT2 cells are present in IPF which are thought to play a central role in the initiation of fibrosis [1], and the regenerative ability of progenitor AT2 cells was proposed to be decreased in IPF [20]. Interestingly, in the present study, IL-11 exposed PCLS showed a decreasing trend in gene expression of alveolar cell types combined with an upregulation of *Foxj1* expression. Though this finding warrants more study, it may relate to bronchiolisation; a pathological process in IPF where alveolar spaces are covered with bronchiolar cell types [1]. Microscopic honeycombing was previously found to be specifically associated with increased expression of cilium-associated genes in a subset of IPF patients [44]. The skewed balance of differentiated alveolar and airway epithelial cell types in the presence of IL-11 could be caused by IL-11 primarily damaging AT2 cells or inhibiting the regenerative response of progenitor AT2 cells. As a result, the alveolar epithelium may not be properly repaired, and other progenitor cells may need to fill the exposed alveolar space with a predominance of bronchiolar cell types. Additionally, IL-11 could affect distal airway progenitors with a regenerative ability of both bronchiolar and alveolar epithelial tissue, by skewing cell fate towards airway cell types, leaving alveolar types insufficiently repopulated.

Myofibroblasts are typically regarded as one of the key cell types in IPF pathogenesis since they are prominent producers of ECM [1]. The effects of IL-11 on fibroblasts have been described previously, where it was found to induce fibroblast to myofibroblast transition and stimulate their deposition of collagen [12]. Surprisingly, here we did not find significant effects on fibrosis marker gene expression in fibroblasts and PCLS exposed to IL-11. It is possible that IL-11 simply does not induce large differences of the expression of fibrosis markers on mRNA level, which was also seen in previous literature [12]. Alternatively, since IL-11 did significantly affect epithelial cells, IL-11 induced epithelial dysfunction may precede the development of fibrosis. Indeed, IL-11 mRNA is also overexpressed in unaffected lung tissue from IPF patients compared to control lung tissue, which suggests IL-11 plays a role in early fibrotic processes [16].

Since lung epithelial progenitor cell behavior is strongly influenced by its microenvironment, we hypothesized that the aberrant epithelial repair response in the presence of IL-11 may be caused by alterations to the progenitor cell niche [45]. Fibroblasts are known to have an important progenitor cell supporting role through paracrine signaling [45], and are additionally present in the organoid co-culture system with epithelial cells. Therefore, we assessed whether IL-11 disturbs fibroblast epithelial cell cross-talk in relation to regeneration. Though IL-11 treatment of primary human fibroblasts resulted in few differentially expressed genes, several of these genes and differentially regulated pathways may relate to fibrosis. While IL-11 induced a down-regulation of *Tgfb1* gene expression in PCLS (figure S4B), and gene expression studies in fibroblasts (figure S6) showed no effect of IL-11 on the expression of *TGFB1*, the *TGFβ pathway* was upregulated in the IL-11 sequencing data set. It is possible that IL-11 stimulates TGFβ signaling using post-transcriptional mechanisms, for example by modulating factors responsible for activating latent TGFβ, thereby promoting pro-fibrotic processes [46]. IL-11 could also upregulate genes that are included in the TGFβ pathway without affecting TGFβ expression itself. In addition to the *TGFβ pathway*, pathways involved with inflammation, such as *TNFα signaling via NF-κB*, *inflammatory response*, *IFNγ response*, *complement* and *IFNα response* were upregulated in fibroblasts exposed to IL-11. IL-11 is known to have immunomodulatory effects [6,7]. However, the role of these upregulated inflammatory pathways [47–55] as well as the role of inflammation in general in IPF is still not completely clear [1]. IL-11 also altered metabolic pathways in primary fibroblasts, such as the *reactive oxygen species (ROS) pathway*. Indeed, mitochondrial ROS was previously found to be increased in IPF tissue and in IPF fibroblasts specifically [56]. *HK2*, *LDHA* and the *glycolysis pathway* were also upregulated, suggesting IL-11 stimulates glycolytic activity in fibroblasts, which was previously found to be increased in IPF tissue. Moreover, glycolytic reprogramming was shown to induce fibroblast to myofibroblast transition, though this was contradicted by other studies [57–59]. Additionally, the *unfolded protein response* (UPR) is upregulated, which can be activated in response to ER stress upon accumulation of misfolded proteins in the ER. ER stress and UPR have been associated with IPF pathogenesis, since they induce fibroblast to myofibroblast transition and the production of collagen by myofibroblasts. The activation of the UPR leads to the degradation of unfolded or misfolded proteins and the general reduction of protein translation with the exception of chaperone and redox proteins, in order to restore cellular homeostasis. The fibroblast secretome may therefore be altered [60].

A downregulated gene of particular interest was *ELN*. In IPF patients, evidence of increased elastin degradation has been reported [61,62]. Since elastin is essential for successful alveologenesis during embryonic development [63] it may also play a role in regenerative mechanisms. Moreover, IL-11-induced myofibroblasts remodel the ECM and increase its stiffness [1], which may affect epithelial cell behavior as well. Finally, *Notch signaling* was downregulated upon IL-11 exposure. Conversely, Notch signaling was found to stimulate myofibroblast differentiation [64], and sustained activation of Notch is related to abnormal epithelial repair with alveolar cyst formation, which is associated with fibrosis [65]. However, Notch is also essential for alveologenesis during embryonic development [66] and for regeneration of club cells after injury to the airway epithelium [67]. Additionally, alveolar repair was found to rely on dynamic Notch signaling, where Notch activity is initially needed to stimulate progenitor cell activation after which Notch signaling is inactivated to support differentiation towards alveolar cell types [65]. Clearly, the role of Notch signaling in epithelial repair is complex and incompletely understood, and its dysregulation may have implications for abnormal re-epithelialization. Overall, the gene expression data show IL-11 exposure in fibroblasts contributes to mechanisms related to pro-fibrotic processes and possibly altered intercellular communication, though the effect on gene expression level is limited.

A limitation of our study is that IL-11 was proposed to act primarily through post-transcriptional mechanisms [12], and its effects may not be evident using only gene expression studies. Indeed, gene expression changes in response to IL-11 were not always apparent in our studies, though it is also possible that the exposure to IL-11 was not long enough to induce noticeable effects. We aimed to explore whether fibroblast-epithelial communication mechanisms were distorted by IL-11, but we were not able to identify many of such mechanisms due to the low response on gene expression of the fibroblasts. Proteomics would be a potential approach to give new insights in these questions. Furthermore, we describe lung structures that show positive immunohistochemical staining of IL-11 and IL-11R, but some cell types such as quiescent fibroblasts and specific immune cells are difficult to recognize without cell-specific markers, and could therefore not be characterized in terms of their expression of IL-11(R). In addition, since IL-11 is a secreted protein, we cannot exclude the possibility that the IL-11 we localized does not solely represent tissues and cells that produce it. However, we noted IL-11 staining in AT2 cells in areas of AT2 cell hyperplasia, in which we did not observe any IL-11R staining, indicating IL-11 positivity likely does not include IL-11Rα-bound IL-11. Finally, the expression and response of IL-11 showed large variability in our studies. Since IPF is known to be more prevalent in men than in women [1], we hypothesized that sex differences could play a role in the response to IL-11. However, our data suggest that the variability of the effect of IL-11 is not due to sex differences in mice. The expression of IL-11 and its receptor in human lung tissue also displayed variability, both intra- and interindividually, which could be related to factors such as age, disease severity, disease activity and smoking status. If IL-11 were to be used as a therapeutic target in the future, the heterogeneity of IL-11 should be addressed to ensure optimal treatment strategy.

In summary, IL-11 disturbs alveolar organoid formation as a model of alveolar regeneration *in vitro*, which may relate to dysfunctional epithelial repair responses in IPF. Though we did not find convincing evidence for a role of IL-11 in disrupting the epithelial progenitor support function of fibroblasts, we show that various other cell types also express IL-11 and IL-11R in the human lung, including epithelial cells, macrophages and SMCs, and their IL-11 driven responses related to fibrosis and dysregulated epithelial repair will be interesting to uncover [12].

## Acknowledgements

We thank Sophie Bos for technical support. The stainings of IL-11 and IL-11R on lung tissue performed in this manuscript were conducted as part of the HOLLAND (HistopathOLogy of Lung Aging aNd COPD) project. The HOLLAND project was initiated and supervised by Corry-Anke Brandsma, Wim Timens, and Janette Burgess, technical support was provided by Marjan Reinders-Luinge, Anja Bakker and Theo Borghuis, and image analyses pipelines were developed by Theo Borghuis, Maunick Lefin Koloko Ngassie and Niek Bekker.

## Supplementary data

Summary

Materials and Methods

Figure S1: the gene expression pattern of IL-11 and IL-11R in the human lung.

Figure S2: IL-11 staining in the human lung is variable amongst individuals, but also within single tissue sections.

Figure S3: inter- and intra-individual variability of IL-11 receptor (IL-11R) staining in the human lung.

Figure S4: fibrosis marker genes are unaffected by IL-11 in lung slices.

Figure S5: recombinant human IL-11 activates downstream signaling pathways JAK/STAT3 and MEK/ERK in CCL206 fibroblasts.

Figure S6: IL-11 does not influence gene expression of established organoid supporting factors in primary human fibroblasts.

References

