## Supplementary information for "IL-11 disrupts alveolar epithelial progenitor function"

Antonius Deusinglaan 1

9713 AV Groningen, the Netherlands

**: current affiliation: Newcells Biotech, Newcastle upon Tyne, United Kingdom

***: authors contributed equally

Key words: lung fibrosis, pulmonary regeneration, interleukin 11, cell communication, lung organoids

**Methods**

Western blot

CCL206 fibroblasts were seeded in a 6-wells plate at a density of 500,000 cells per well and allowed to settle for 24 hours, after which they were serum-starved for 24 hours with 0.5% FBS in DMEM:Ham’s F12 (1:1) supplemented with 2 mM L-glutamine, 100 U/ml penicillin/streptomycin, and 1% amphotericin B. They were treated with 1-100 ng/ml recombinant mouse IL-11 (Peprotech, Cranbury, NJ, USA, #220-11) or rhIL-11 (R&D systems, #218-IL) for 30 minutes, then washed with ice-cold PBS and lysed in modified RIPA buffer with protease inhibitors aprotinin, leupeptin, and pepstatin, and phosphatase inhibitors β-glycerolphosphate (Ser/Thr phosphatase inhibitor), sodium orthovanadate (Tyr phosphatase inhibitor) and sodium fluoride (Ser/Thr phosphatase inhibitor), and finally subjected to sonification. 30 µg of protein and 5 µl PageRuler Plus Protein Ladder (Thermo Scientific, #26620) were loaded on a 10% SDS-PAGE gel and run using 1x ELFO (0.025 M Tris, 0.25 M glycine and 0.1% SDS in UP water). Proteins were transferred to a nitrocellulose membrane using transfer buffer (0.025 M Tris, 0.192 M glycine, 0.1% SDS and 20% methanol (v/v) in UP), after which the membrane was cut between the 70 and 55 kDa markers. Membranes were blocked for 1.5 hours with 1x ROTI block (Carl Roth, Karlsruhe, Germany, #A151.2) for α-tubulin, ERK and p-ERK, 5% milk for STAT3 and 5% BSA for p-STAT3. Membranes were incubated with the primary antibodies mouse α-tubulin (final concentration 0.5 µg/ml, Sigma-Aldrich, #T6074, RRID: AB_477582), rabbit ERK1/2 (1:1000, Cell Signaling, Danvers, MA, USA, #9102, RRID: AB_330744), rabbit p-ERK1/2 (1:1000, Cell Signaling, #9101, RRID: AB_331646), mouse STAT3 (1:1000, Cell Signaling, #9139, RRID: AB_331757) and rabbit p-STAT3 (1:2000, Cell Signaling, #9145, RRID: AB_2491009) overnight at 4 °C with rocking. Subsequently, membranes were incubated with secondary peroxidase antibodies goat anti-rabbit (final concentration 0.33 µg/ml, Sigma-Aldrich, #12-348, RRID: AB_390191) or rabbit anti-mouse (1:3000, Sigma-Aldrich, #A9044, RRID: AB_258431) for 2 hours at room temperature with rocking. Protein bands were visualized using ECL Western Blotting Substrate in a GBox iChemi XR system (Syngene, Bangalore, India) using GeneSnap ver 7.12 and quantified using GeneTools ver 4.01 software. Band intensity of the protein of interest was normalized to the intensity of α-tubulin.

***Figure S1: the gene expression pattern of IL-11 and IL-11R in the human lung.*** *RNA sequencing data of the human lung adapted from the IPF cell atlas [1,2], showing the expression of IL-11 (A) and IL-11 receptor (IL-11R) (B) in the various cell types of the lung.*

***Figure S2:*** ***IL-11 staining in the human lung is variable amongst individuals, but also within single tissue sections.*** *IL-11 immunohistochemistry was performed on human lung tissue sections of control and idiopathic pulmonary fibrosis (IPF) donors, which was visualized with NovaRED (red). Sections were counterstained with hematoxylin (blue). Labels show whether the staining was relatively low, medium (med), or high. Black arrows indicate positively stained structures, whereas yellow arrows indicate corresponding structures in the same tissue section that appear negative or stained to a lesser extent. (A) shows the negative control. Example images (B-E) are from the control group, whereas (F-I) are from IPF donors. All scale bars in the overview images are 1 mm, scale bars in enlarged images are 500 μm.*

***Figure S3: inter- and intra-individual variability of IL-11 receptor (IL-11R) staining in the human lung.*** *Human lung tissue of control and idiopathic pulmonary fibrosis (IPF) donors were stained for IL-11R, developed with NovaRED (red), and counterstained with hematoxylin (blue). Labels indicate relative level of staining; low, medium (med), or high, compared to other sections. Black arrows show structures positive for IL-11R, and yellow arrows signal the same structures within the tissue section that are negative or stained to a lesser extent. (A) shows the negative control. Representative images (B-E) belong to the control group, and (F-I) are from IPF donors. The scale bars in overview images are 1 mm, in magnified images they are 500 μm.*

***Figure S4: fibrosis marker genes are unaffected by IL-11 in lung slices.*** *Precision cut lung slices (PCLS) of wild type mice were treated with 100 ng/ml rhIL-11 for 48 hours, after which RNA was isolated from whole slices and gene expression studies were performed using PCR. (A) Gene expression of club cell marker* (Scgb1a1) *and goblet cell marker* (Muc5ac)*. (B) Expression of fibrosis-associated genes collagen 1* (Col1a1), *fibronectin 1* (Fn1), *fibulin 1* (Fbln1)*, MMP7* (Mmp7) *and TGFβ* (Tgfb1)*. (C) IL-11* (Il11) *and IL-11 receptor* (Il11ra1) *gene expression*. (A-C) *N=9 for* Fbln1, *N=10 for all other genes. For all genes: paired samples Wilcoxon test (median is shown).*

***Figure S5: recombinant human IL-11 activates downstream signaling pathways JAK/STAT3 and MEK/ERK in CCL206 fibroblasts.*** *CCL206 mouse lung fibroblasts were exposed to 1-100 ng/ml rhIL-11 and rmIL-11 for 30 minutes, after which they were lysed in modified RIPA buffer and used for Western Blot experiments of STAT3 and ERK1/2 activation. (A) Representative uncropped western blots of untreated CCL206 cells (Un), CCL206 cells exposed to vehicle (C) and CCL206 cells in response to 1-100 ng/ml IL-11. Protein bands of total STAT3 and ERK1/2, their phosphorylated forms, and respective α-tubulin loading controls are shown. Samples were loaded onto 2 gels, of which 1 was used for STAT3 and ERK1/2, and the other for p-STAT3 and p-ERK1/2. Membranes were cut between the 70 and 55 kDa marker, after which the 250-70 kDa membrane was incubated with STAT3 or p-STAT3 antibody, and the 55-10 kDa membrane was incubated with ERK1/2 or p-ERK1/2 antibody and subsequently reprobed for α-tubulin. (B) Quantification of the phosphorylation of STAT3 (N=3) in response to rhIL-11 where the p-STAT3/STAT3 band intensity ratio (corrected for α-tubulin band intensity) is shown. (C) Quantification of the phosphorylation of ERK1/2 (N=3) after rhIL-11 exposure, showing the band intensity ratio of p-ERK1/2/ERK1/2 (corrected for α-tubulin band intensity).*

***Figure S6****:* ***IL-11 does not influence gene expression of established organoid supporting factors in primary human fibroblasts.*** *Primary human fibroblasts of either control or idiopathic pulmonary fibrosis (IPF) donors were exposed to 100 ng/ml rhIL-11 for 24 hours, after which RNA was isolated and PCR studies were performed. (A) Gene expression of the epithelial cell supporting factors* FGF2*,* FGF7*,* FGF10*, hepatocyte growth factor* (HGF) *and* AXIN2 *(Wnt signaling). (B) Expression of fibrosis marker genes fibronectin 1* (FN1) *and TGFβ* (TGFB1)*. (C) Gene expression of IL-11 related genes IL-11* (IL11) *and IL-11R* (IL11RA). *(A-C) Control donors (N=6), IPF donors (N=5), Two-way ANOVA with Sidak’s post hoc test on log transformed data.*
