## Supplementary figures and images for "IL-11 disrupts alveolar epithelial progenitor function"

### Figure S1

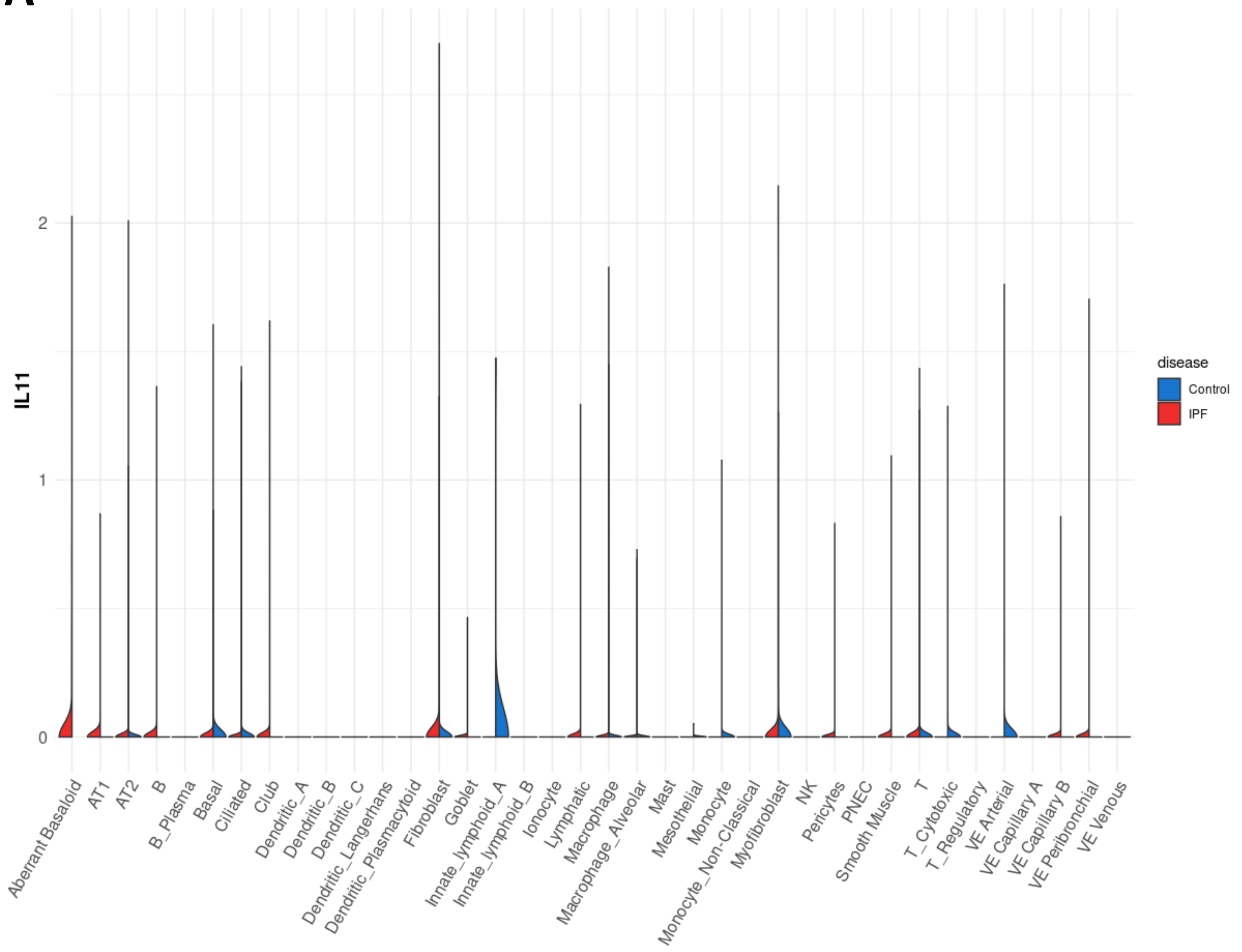

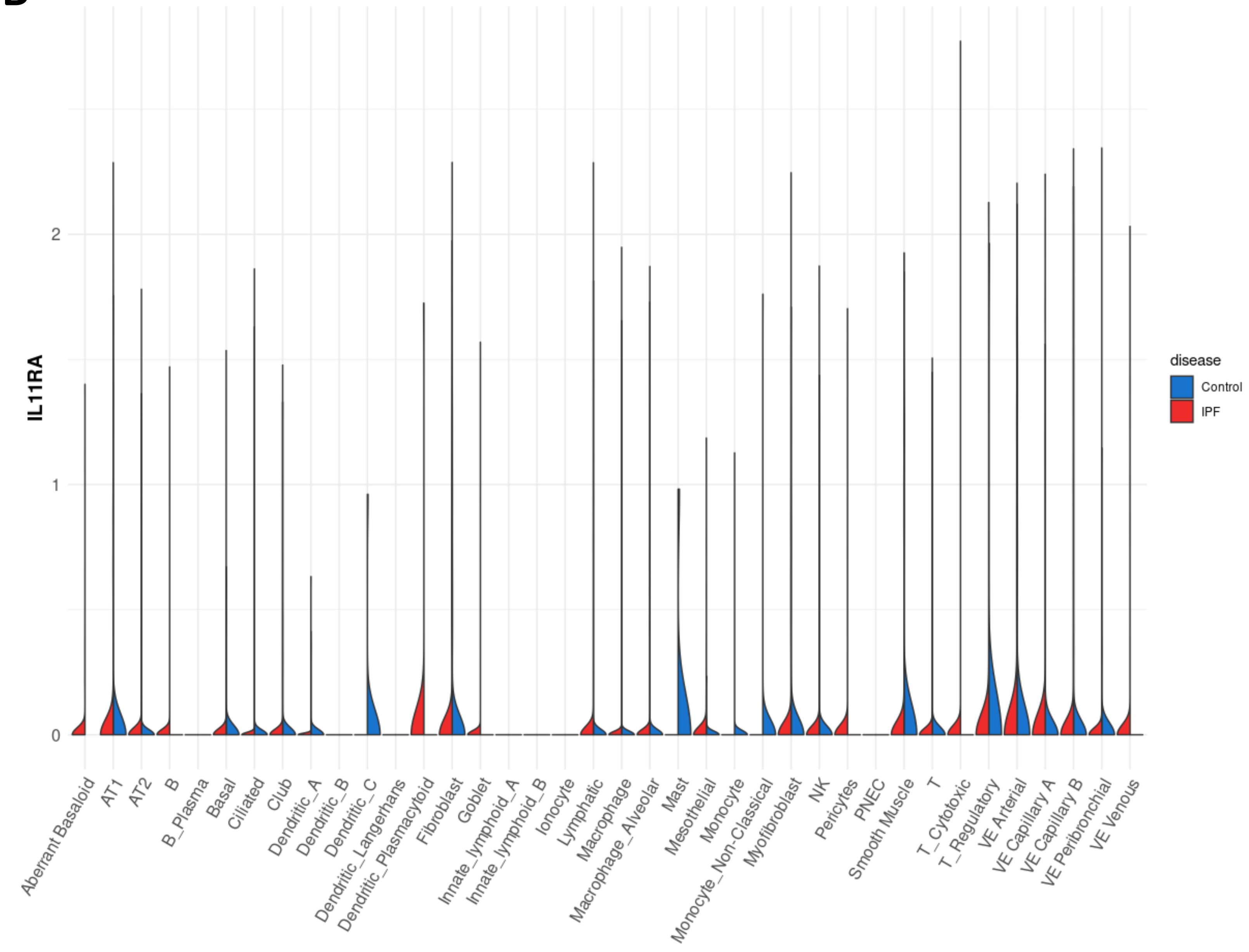

### Figure S2

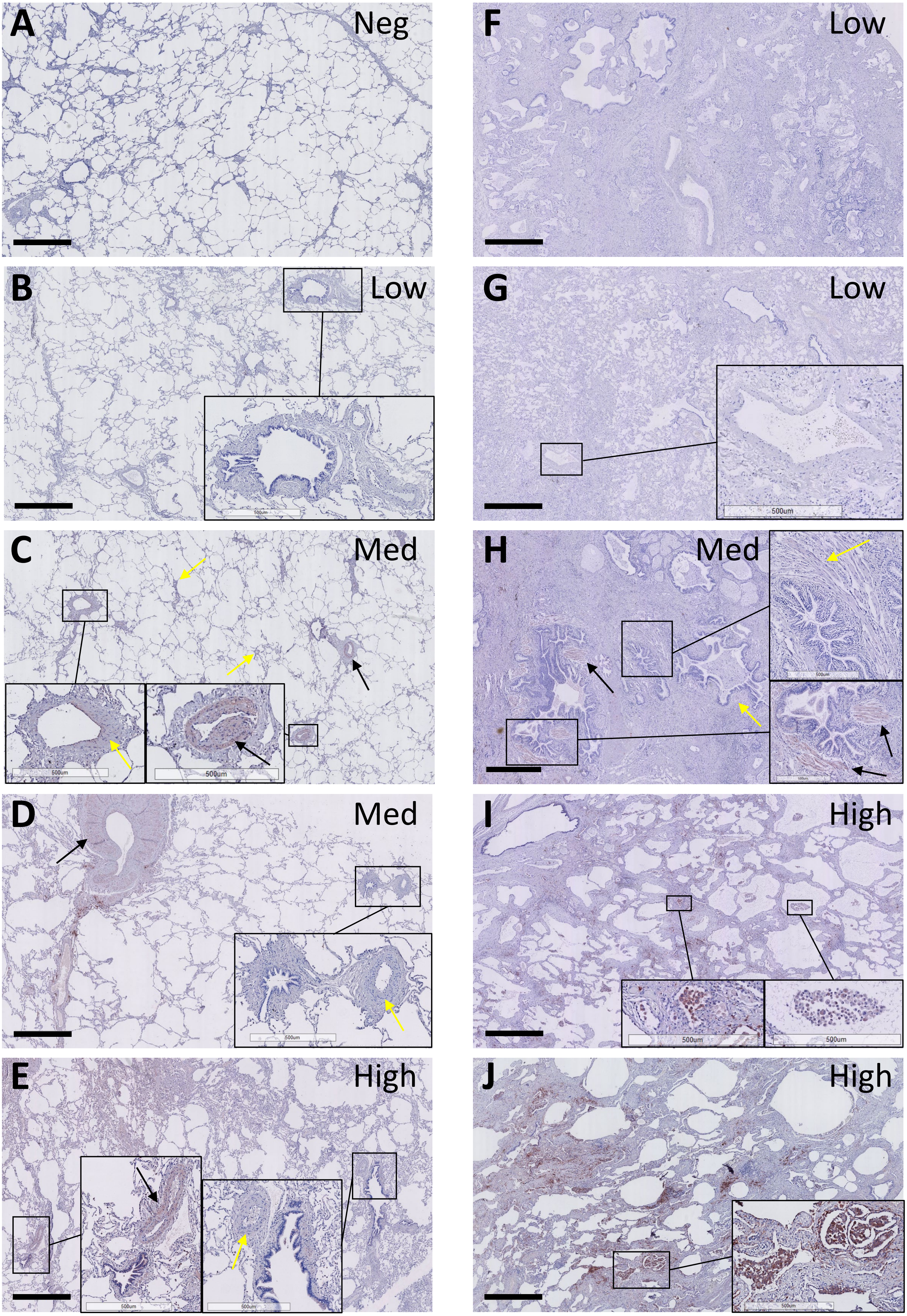

### Figure S3

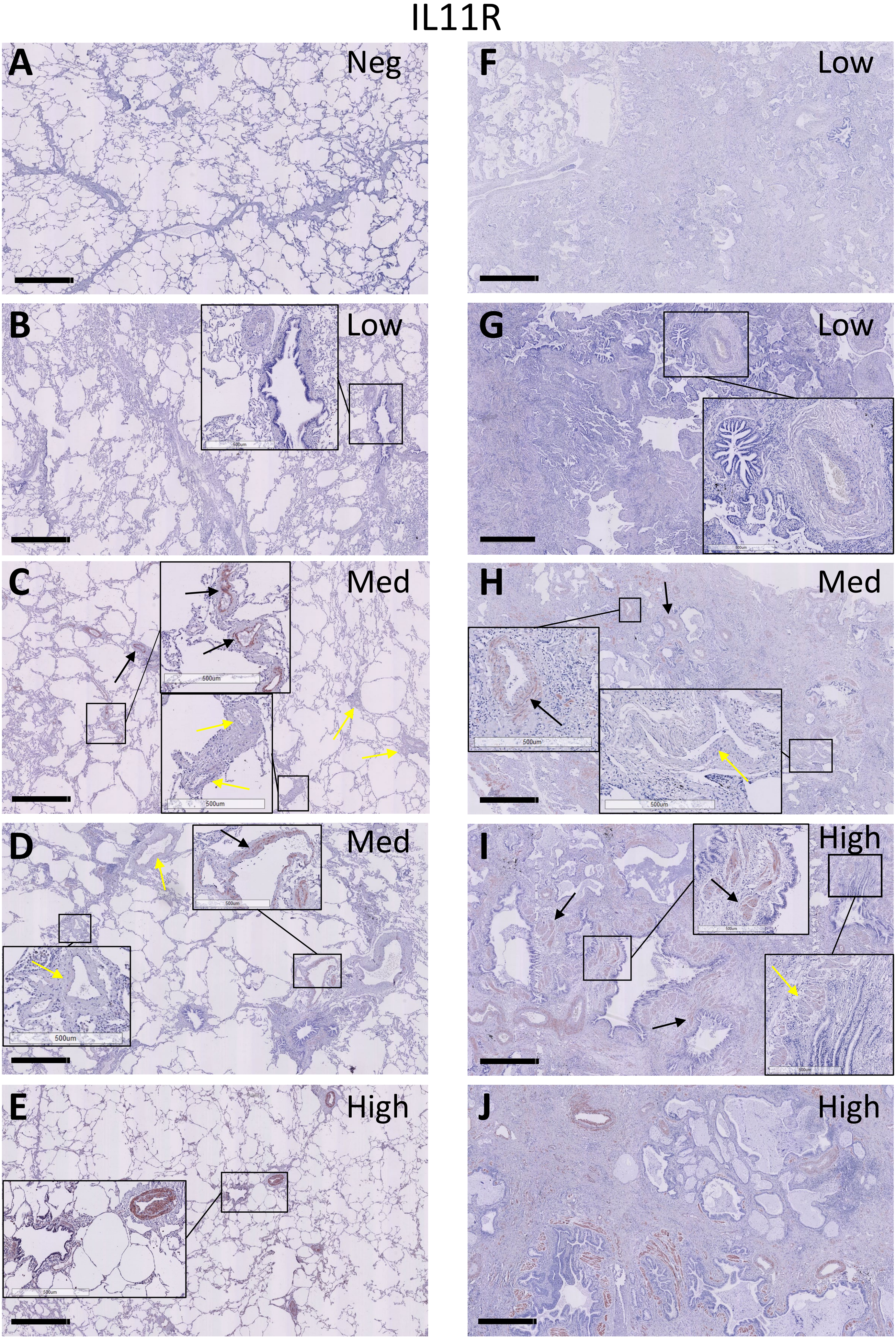

### Figure S4

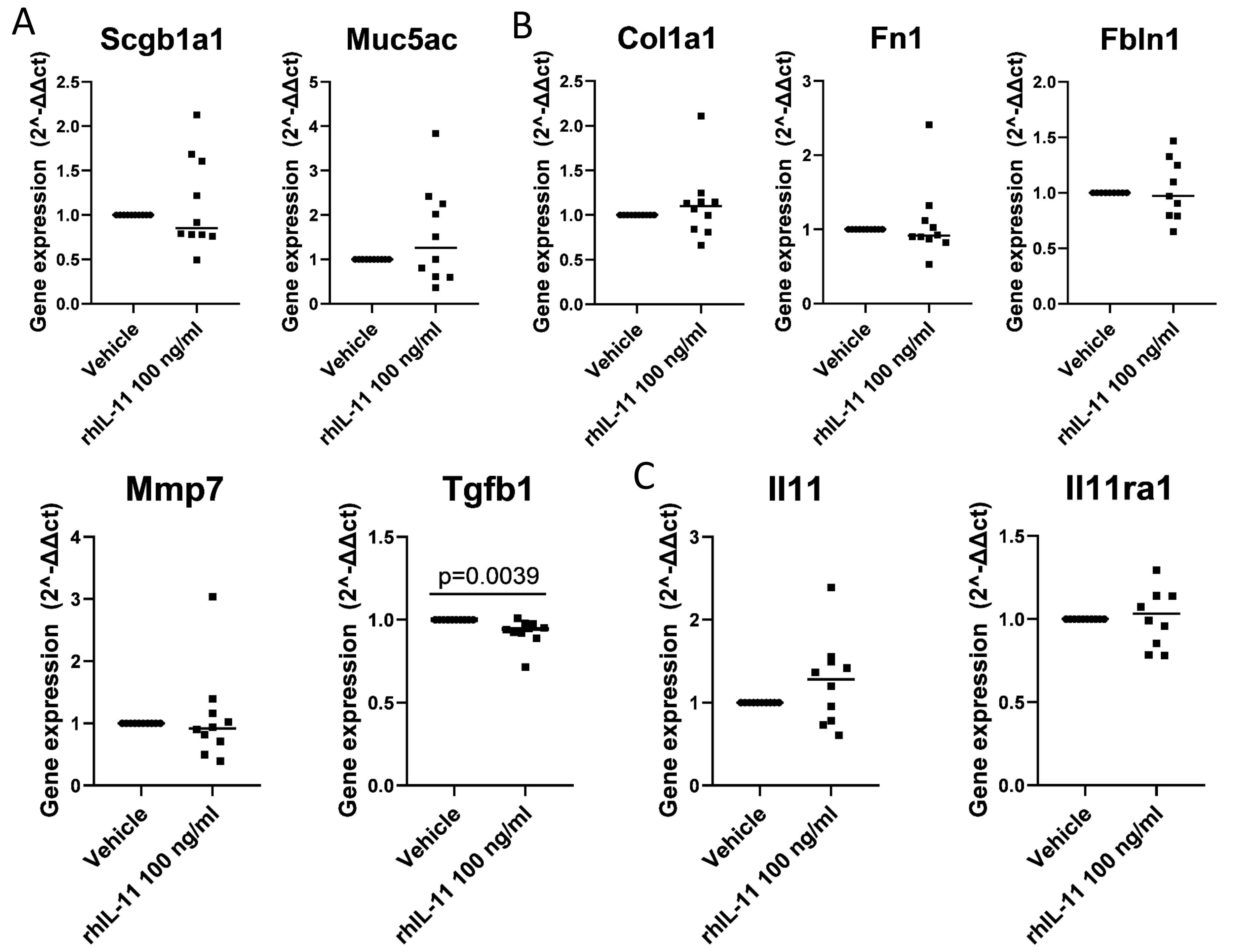

### Figure S5

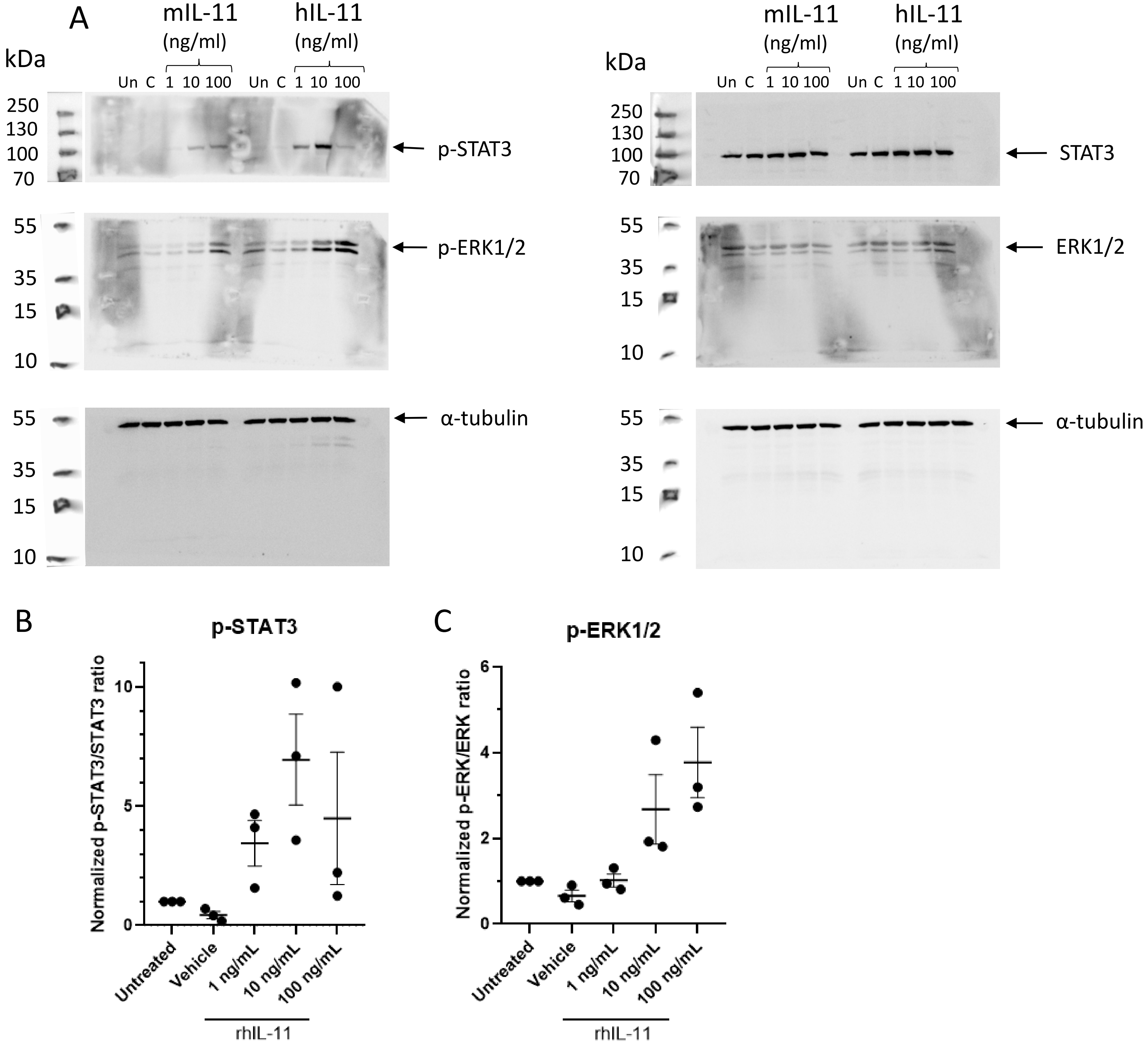

### Figure S6

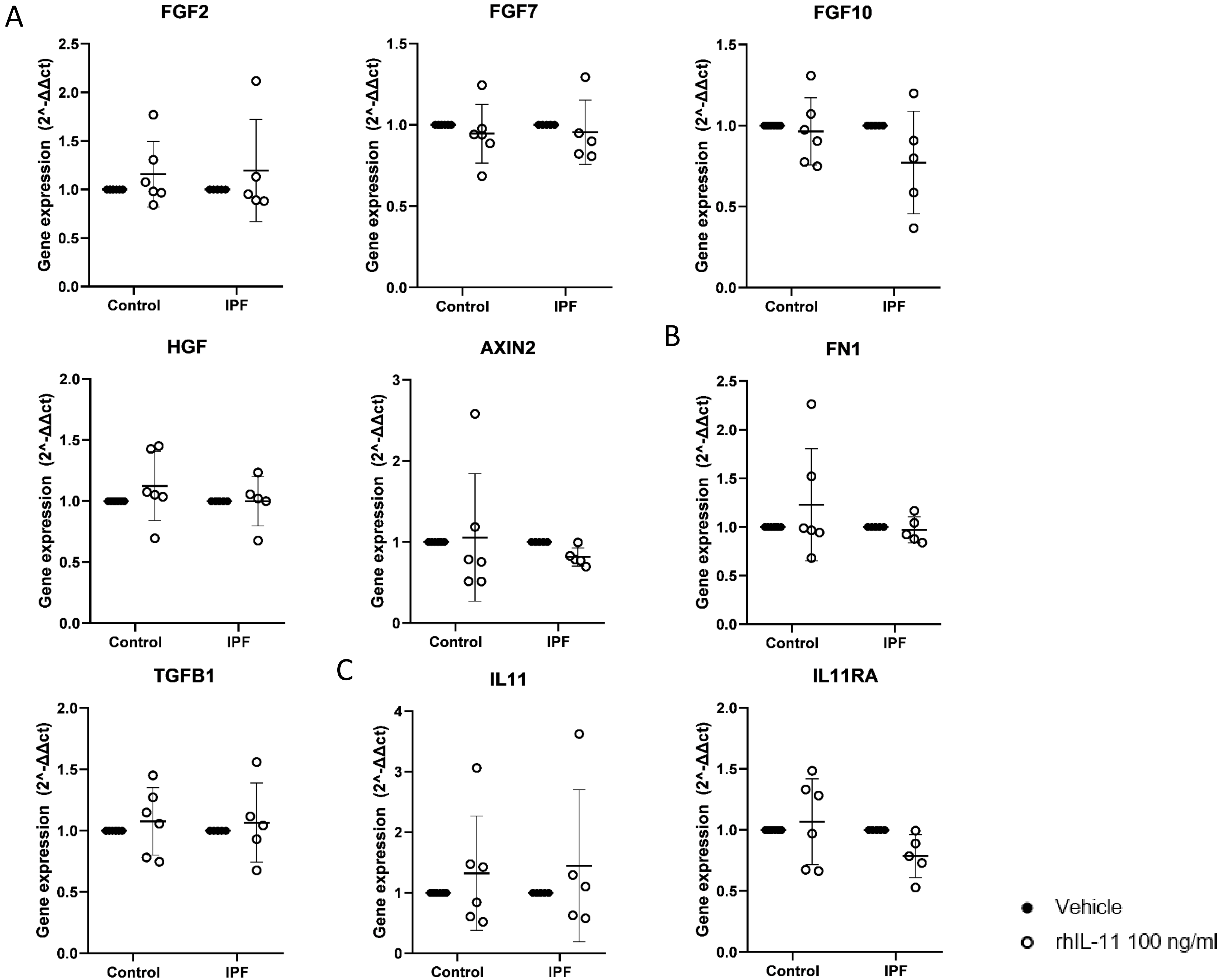
