## Supplementary material for "IL-11 disrupts alveolar epithelial progenitor function": Table S1

Table S1: primer sequences used for analyses on human samples

| **Gene** |  | **Sequence (5’-3’)** |
| --- | --- | --- |
| *B2M* | Fw | TGG AGG CTA TCC AGC GTA CT |
|  | Rv | CGG ATG GAT GAA ACC CAG ACA |
| *SDHA* | Fw | GCA TGC CAG GGA AGA CTA CA |
|  | Rv | ACG GGT CTA TAT TCC AGA GTG AC |
| *HMBS* | Fw | TGGACCTGGTTGTTCACTCCTT |
|  | Rv | CAACAGCATCATGAGGGTTTTC |
| *FGF2* | Fw | AAA AAC GGG GGC TTC TTC CT |
|  | Rv | TGT AGC TTG ATG TGA GGG TCG |
| *FGF7* | Fw | CCC TGA GCG ACA CAC AAG A |
|  | Rv | CCA CAA TTC CAA CTG CCA CTG |
| *FGF10* | Fw | ATG TCC GCT GGA GAA AGC TA |
|  | Rv | CCC CTT CTT GTT CAT GGC TA |
| *HGF* | Fw | CTG GTT CCC CTT CAA TAG CA |
|  | Rv | CTC CAG GGC TGA CAT TTG AT |
| *AXIN2* | Fw | TGT GAG GTC CAC GGA AAC TG |
|  | Rv | CTG CCC ACA CGA TAA GGA GG |
| *FN1* | Fw | AAT GCA CCA CAG CCA TCT CA |
|  | Rv | GTC ACT TCT TGG TGG CCG TA |
| *TGFB1* | Fw | GTA CCT GAA CCC GTG TTG CT |
|  | Rv | GAA CCC GTT GAT GTC CAC TT |
| *IL11* | Fw | GAG TTT CCC CAG ACC CTC GG |
|  | Rv | GTA GGA CAG TAG GTC CGC TC |
| *IL11RA* | Fw | CCA GCC AGA TCA GCG GTT TA |
|  | Rv | CCA GTG GGT TCA CCT CAG TC |
