## Supplementary material for "IL-11 disrupts alveolar epithelial progenitor function": Table S2

Table S2: primer sequences used for analyses on mouse samples

| **Gene** |  | **Sequence (5’-3’)** |
| --- | --- | --- |
| *Rpl13a* | Fw | AGA AGC AGA TCT TGA GGT TAC GG |
|  | Rv | GTT CAC ACC AGG AGT CCG TT |
| *B2m* | Fw | ATG GGA AGC CGA ACA TAC TG |
|  | Rv | CAG TCT CAG TGG GGG TGA AT |
| *Actb* | Fw | ATC GTG CGT GAC ATC AAA GA |
|  | Rv | ATG CCA CAG GAT TCC ATA CC |
| *Hopx* | Fw | CGACTTTCAGTGGTTCCTGC |
|  | Rv | GTGTGGAAGTCTGGGCGAG |
| *Sftpc* | Fw | GGAGCACCGGAAACTCAGAA |
|  | Rv | GGAGCCGCTGGTAGTCATAC |
| *Scgb1a1* | Fw | GGCCCTCCTCATGGAATCAG |
|  | Rv | GCATTTTGCAGGTCTGAGCC |
| *Muc5ac* | Fw | GAG ATG GAG GAT CTGG |
|  | Rv | GCA GAA GCA GGG AGT GGT AG |
| *Foxj1* | Fw | CGG CCA TCT ACA AGT GGA TCA |
|  | Rv | CTT GAA GGC CCC ACT GAG CA |
| *Fn1* | Fw | ACCACCCAGAACTACGATGC |
|  | Rv | GGAACGTGTCGTTCACATTG |
| *Col1a1* | Fw | CACCCTCAAGAGCCTGAGTC |
|  | Rv | GTTCGGGCTGATGTACCAGT |
| *Fbln1* | Fw | AGAACTATCGCCGCTCCGCA |
|  | Rv | CCACCGCTGGCACTTGGATG |
| *Mmp7* | Fw | GGT GTG GAG TGC CAG ATG TT |
|  | Rv | TAT CCG CAG TCC CCC CAA CTA |
| *Tgfb1* | Fw | GGACTCTCCACCTGCAAGAC |
|  | Rv | GACTGGCGAGCCTTAGTTTG |
| *Il11* | Fw | TGT TCT CCT AAC CCG ATC CCT |
|  | Rv | CAG GAA GCT GCA AAG ATC CCA |
| *Il11ra1* | Fw | ATC CGT ACC TGG TTA CCC GA |
|  | Rv | CCC AGC CAC AGC ATC TGT TA |
